# mEnrich-seq: Methylation-guided enrichment sequencing of bacterial taxa of interest from microbiome

**DOI:** 10.1101/2022.11.07.515285

**Authors:** Lei Cao, Yimeng Kong, Yu Fan, Mi Ni, Alan Tourancheau, Magdalena Ksiezarek, Edward A. Mead, Tonny Koo, Melissa Gitman, Xue-Song Zhang, Gang Fang

**Affiliations:** Department of Genetics and Genomic Sciences, Icahn School of Medicine at Mount Sinai, New York, NY 10029, USA; Department of Pathology, Molecular and Cell-based Medicine, Icahn School of Medicine at Mount Sinai, New York, NY 10029, USA; Center for Advanced Biotechnology and Medicine, Rutgers University, New Brunswick, NJ 08901, USA

## Abstract

Metagenomics has enabled the comprehensive study of microbiomes. However, many applications would benefit from a method that can sequence specific bacterial taxa of interest (pathogens, beneficial microbes, or low-abundance taxa), but not the vast background of other taxa in a microbiome sample. To address this need, we developed mEnrich-seq, a method that can enrich taxa of interest from metagenomic DNA before sequencing. The core idea is to exploit the self vs. non-self genome differentiation provided by natural bacterial DNA methylation and rationally choose methylation-sensitive restriction enzymes (REs), individually or in combination, to deplete host DNA and most background microbial DNA while enriching bacterial taxa of interest. This core idea is integrated with library preparation procedures in a way that only non-digested DNA libraries are sequenced. We performed in-depth evaluations of mEnrich-seq and demonstrated its use in several applications to enrich (up to 117-fold) genomic DNA of pathogenic or beneficial bacteria from human urine and fecal samples, including several species that are hard to culture or of low abundance. We also assessed the broad applicability of mEnrich-seq and found that 3130 (68.03%) of the 4601 strains with mapped methylomes to date can be targeted by at least one commercially available RE, representing 54.78% of the species examined in this analysis. mEnrich-seq provides microbiome researchers with a versatile and cost-effective approach for selective sequencing of diverse taxa of interest directly from the microbiome.

## Introduction

Microbiomes play important roles in human health, diseases, and drug responses^1–3^ . In addition, human and environmental microbiomes are rich resources to discover genes and molecules with biomedical, bioengineering, and industrial impacts^4–9^. Owing to a number of advanced metagenomic tools^10–15^, many studies have reported that the dysbiosis of microbiota, or the loss of microbial diversity, is not only associated with a higher risk of proliferating pathogenic microbes but is linked to many complex diseases^16–19^. In many applications, researchers hope to specifically examine certain bacterial taxa of interest in a microbiome sample, rather than sequencing all the species^20–24^. For example, a few enteric pathogens are of particular interest in infectious diseases^23, 24^, while certain commensal bacteria are the target of discovery in terms of microbe-host interaction, immune homeostasis, or biosynthetic gene clusters^7–9, 20–22^.

Significant challenges arise in the sequencing of bacterial taxa of interest from complex microbiome samples, mostly stemming from the presence of bacterial species with complex genomes, the highly skewed abundance distribution, mixed with viruses, fungi, and mammalian cells^25–27^ . The sequencing throughput is largely consumed by abundant microbes while low-abundant ones cannot be sequenced and studied in a cost-effective manner^28–30^. Although culture can be used to isolate specific bacteria, it is time- and resource-consuming, hard to scale, and sometimes very challenging for certain bacterial taxa that are hard to culture or impossible to culture yet^31–33^. A few technologies have been developed to address this challenge. Host genomic DNA (gDNA) can be eliminated using chemicals such as saponin to selectively lyse mammalian cells, which enables the efficient sequencing of pathogens in upper respiratory tract infection^34, 35^, where biospecimen are dominated by host gDNA. Host genetic material can also be filtered using DNA binding proteins that specifically recognize 5-methylcytosine (5mC), abundant in mammalian genomes^36^. Flow cytometry has also been used to segregate complex microbiomes into small subsets, which can be sequenced and computationally analyzed separately^37^. However, it is challenging for these approaches to enrich specific taxa of interest, taxa with low abundance, or resolve bacterial species and strains with highly similar genomes^37^. Adaptive sampling in the Nanopore platform can examine the first few hundred base pairs^38^, during real-time sequencing, against a collection of reference sequences and to reject the DNA molecules that meet certain criteria, *e.g.,* host DNA or a highly abundant bacterium for which enough data have been already collected^30, 39, 40^. This strategy by adaptive sampling can reduce further consumption of sequencing yield by high abundant species; however, its enrichment efficiency for specific taxa of interest is relatively modest, and importantly insufficient to differentiate between bacterial taxa with highly similar genomes in complex microbiome samples^41, 42^ . Due to the limitations of these existing methods, our capability to examine the genomes of specific bacterial taxa of interest directly from metagenomic DNA is still limited.

To address the need for effective enrichment of bacterial taxa of interest directly from metagenomic DNA, we developed a method taking advantage of bacterial DNA methylation in microbiomes, namely *bacterial epigenome*^43, 44^. In the bacterial kingdom, there are three major forms of DNA methylation: N6-methyladenine (6mA), N4-methylcytosine (4mC) and 5mC, which are catalyzed by methyltransferases (MTases) that apply methyl groups in a highly sequence-specific manner^45^, *i.e.,* some sequence motifs are nearly 100% methylated, while most of the genome is not methylated. The bacterial methylome has three fundamental properties. First, DNA methylation is present in nearly all bacteria (>95%, three methylation motifs in each organism, on average)^44, 46, 47^. Second, all genetic contents (chromosomes, plasmids, and phages) in a single strain share the same set of methylation motifs, and most methylation sites are highly stable over time and conditions^43–45, 48, 49^. Third, since the presence/absence of a MTase is mainly driven by horizontal gene transfer, the methylation motifs are highly variable between different species and different strains of the same species^47, 50, 51^. Based on these three properties, bacterial DNA methylation naturally differentiates self from non-self DNA, which serves as the foundation of restriction modification (R-M) systems, and has been exploited as natural epigenetic barcodes to group highly similar species and strains in metagenomic analyses, namely *methylation binning*^48, 49^.

In this work, our method exploits bacterial DNA methylation motifs to differentiate bacterial taxa from each other *before* sequencing, in order to enrich bacteria of interest and deplete background DNA. The core idea is to rationally choose individual, or combinations of, methylation-sensitive restriction enzymes (REs) that digest host genome and background microbial DNA at non-methylated sequence recognition sites to eliminate the vast background of metagenomic DNA while leaving the gDNA from bacterial taxa of interest intact. This core idea is integrated with library preparation procedures in a way that only non-digested DNA libraries are sequenced, which provides an effective approach for more efficiently allocating sequencing throughput to examining bacterial taxa of interest from the microbiome. We will first describe the method design and the evaluation, and then provide a few applications to demonstrate its utilities in microbiome research. We also assessed the broad utility of mEnrich-seq in a meta-analysis of the 4,601 bacterial methylomes mapped to date and found that 68.03% of bacterial strains can be targeted by mEnrich-seq with at least one commercially available RE, with broad taxonomic diversity spanning 54.78% of the species examined in this analysis.

## Results

### Methylation-guided enrichment for bacterial taxa of interest from metagenomic DNA

The core idea of enriching the gDNA of bacterial taxa of interest from the microbiome is to distinguish gDNA from different taxa based on their distinct characteristics. Here we describe methylation-guided enrichment sequencing of bacterial taxa of interest from microbiomes, namely mEnrich-seq (where ‘m’ stands for methylation, Fig. 1a). For a bacterial taxon of interest, based on its methylation motifs, a methylation-sensitive restriction enzyme with the same recognition motif is chosen to cut the non-methylated motif sites in most of the “background” DNA (Fig. 1a). In contrast, the gDNA of the bacterial taxa of interest (including its chromosome and mobile genetic elements, *e.g*., plasmids) is left intact, enriched over the background, which can be sequenced by various sequencing platforms (Fig. 1a). More specifically, we first use g-TUBEs to shear the DNA size to around 10kb, which facilitate the second step where DNA is ligated to barcode adapters. In the third step, we use the cognate RE corresponding to the methylation motif in the taxon of interest, to cut the background DNA libraries into smaller fragments, while preserving the targeted DNA libraries as the long fragments. Because bacterial DNA methylation motifs usually have 2 to 8 non-degenerate bases (*e.g*. GANTC has four non-degenerate bases; N = A, C, G or T)^45, 47–49^, the choice of fragmentation size in the shearing step (generating 5∼10kb fragments) increases the chance that most gDNA molecules have at least one non-methylated RE site that can be cut: 5∼10kb fragments expected to have 20∼40 sites of a 4-mer motif, 5∼10 sites of a 5-mer motif, and 1∼2.5 sites of a 6-mer motif. In the fourth step, we perform a gel-based size selection, which largely remove the digested shorter DNA fragments. In the fifth step, we use the barcoded primers anchoring the PCR adapter to perform amplification of the long target DNA fragments that have been protected from RE digestion (methylated at restriction sites). In the sixth and final step, the amplified dsDNA is sequenced and analyzed using various metagenomic analysis tools. Because we designed the DNA-to-adapter ligation (step 2) before RE cutting (step 3), only non-digested DNA libraries can be sequenced. This design minimizes allocation of sequencing throughput to background DNA fragments that are still long after RE digestion due to sparse distribution of the RE recognition motif(s) in certain genomes or genomic regions.

**Fig. 1.**
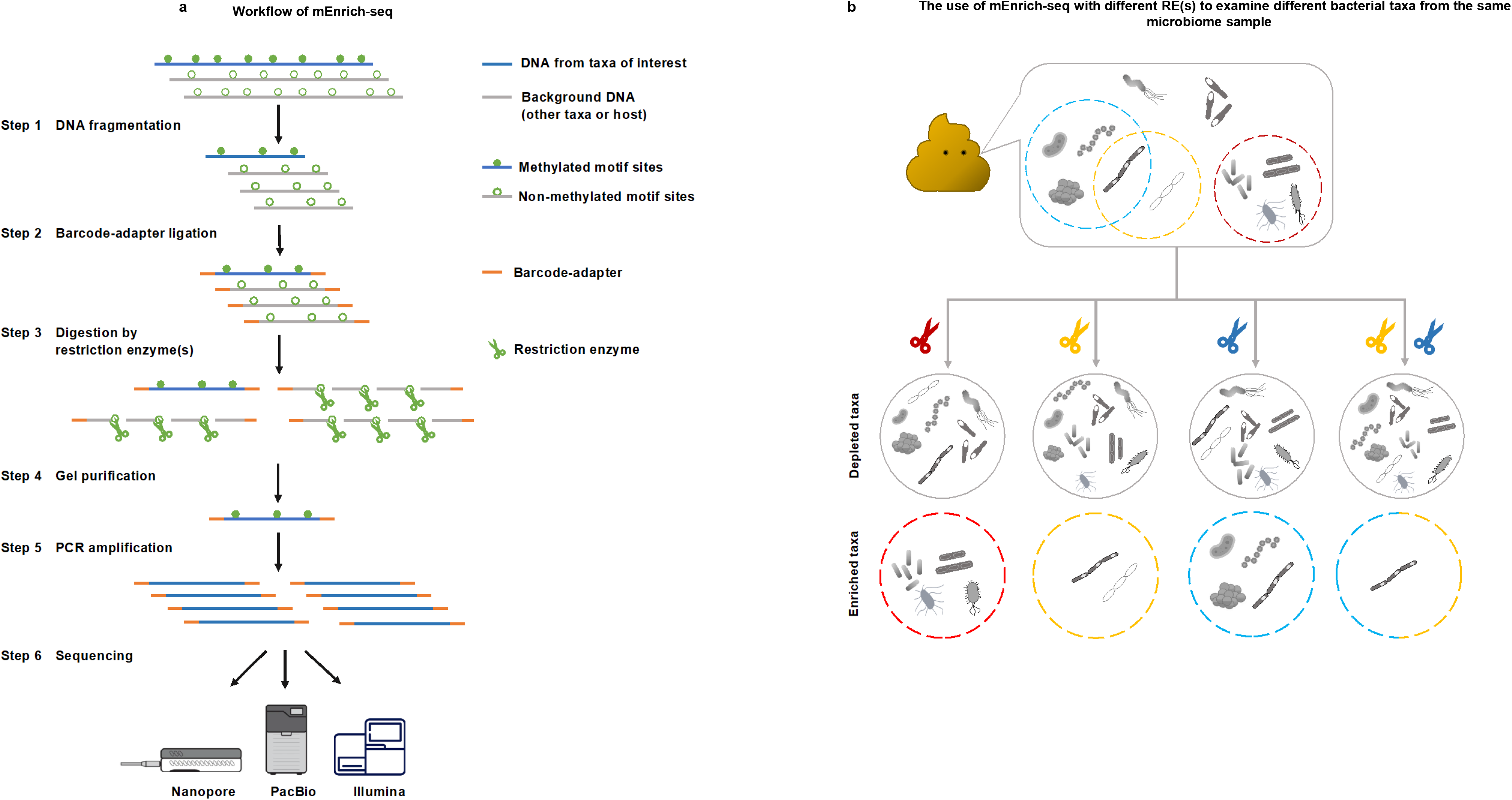
mEnrich-seq workflow and its application. **a,** The mEnrich-seq workflow. **1st**, a metagenomic DNA sample is sheared with a g-TUBE to a selected fragment size. **2nd**, the sheared DNA molecules are end-prepared and ligated to the barcode-adapter. **3rd**, the barcode-adapter-ligated DNA molecules are digested with one or more restriction enzymes (REs, green scissors) that cut the non-methylated motif sites (green circles), leaving the methylated motifs (filled green dots) intact. **4th**, gel-based purification is performed to recover undigested DNA molecules at a high molecular weight. **5th**, purified DNA from the previous step is amplified with PCR**. In the 6th step**, amplified DNA molecules are compatible with multiple sequencing platforms after library preparation per the sequencing platforms’ instructions. **b,** The use of mEnrich-seq with different RE(s) to examine different bacterial taxa from the same microbiome sample. The colored circles with dashed lines highlight bacteria in a microbiome sample that share common methylated motifs, which can be targeted by different REs (scissors with colors corresponding to dashed circles). Each bacterial taxon can have multiple methylation motifs, and thus can be shared between different dashed circles. When we add one RE (*e.g.,* the red scissors represent one RE) to the metagenomic DNA sample, we can enrich a subgroup of bacteria of interest (circled by the red dashed line), by depleting the rest of the bacteria (the gray circle). When applying the individual REs to the microbiome sample separately (represented by the individual yellow and blue scissors), mEnrich-seq allow researchers to examine different subgroups of bacteria in the microbiome sample (circled by individual yellow or blue dashed lines). mEnrich-seq can also be tailored with the use of multiple REs to more specifically enrich bacterial taxa with multiple methylation motifs (as illustrated by both the yellow and blue scissors, enriching the bacterium shared between blue and yellow dashed line).

mEnrich-seq is versatile along three dimensions. First, while this work used Nanopore sequencing, the mEnrich-seq protocol described above is compatible with all sequencing platforms (such as PacBio and Illumina sequencing platforms) by adjusting the insert size (after step 5) and chemistry according to different platforms (see Methods) (Fig. 1a). Second, because bacterial genomes usually have multiple methylation motifs (three on average, over twenty in certain strains)^45, 47–49^, multiple restriction enzymes can be used simultaneously in the third step of mEnrich-seq described above, which can further enhance the enrichment efficiency (see Methods and later sections) (Fig. 1b). Third, some methylation motifs are conserved across different strains of the same species, or across different genera (*e.g.,* G<u>6mA</u>TC in most Gammaproteobacteria)^47, 52^. Therefore, the choice of RE(s) can be tailored to enrich bacterial taxa at different taxonomic levels, which can be helpful depending on specific research goals (Fig. 1b).

We first applied our new method to a mock sample for a proof of concept and illustration. Genomic DNA from *Escherichia coli* (representing bacterial taxa of interest, ∼1.00% by mass), human Lymphoblastoid Cell Line (LCL) (representing host gDNA, >94.00% by mass), and *Neisseria gonorrhoeae* (representing background bacterial taxa of no interest, ∼5.00% by mass) were mixed together (two replicates). Because GATC sites are methylated (6mA) in the *E. coli* genome^43, 44, 46, 53^ (a site occurs approximately every 250 bp) but not in *N. gonorrhoeae* or the human genome, the methylation-sensitive RE DpnII (which cut dsDNA at non-methylated GATC sites, but sensitive to methylated G<u>6mA</u>TC) was chosen for mEnrich-seq to selectively enrich *E. coli* gDNA while digesting *N. gonorrhoeae* (only 35 of the 2434 GATC sites are methylated owing to the overlapping with GGTG<u>6mA</u> methylation motif) and human gDNA with hardly detectable methylated GATC sites. After digesting the DNA mock samples with DpnII, we prepared the library for Nanopore long-read sequencing and generated 200Mb of data (mean read length of ∼5kb; Supplementary Table 1). For comparison, we also sequenced the mock samples without mEnrich-seq protocol, labeled as “input” (standard metagenomics).

Compared with the input data, mEnrich-seq greatly increased the proportion (%) of reads mapped to the *E. coli* genome from 0.91% to 70.75% and 1.20% to 71.41% in the two mock replicates, respectively (Fig. 2a), ∼70-fold enrichment (Fig. 2b), illustrating the efficient enrichment of gDNA of bacteria of interest. mEnrich-seq reads mapping showed that the *E. coli* genome can be covered without systematic bias for both mock samples (>99.96% of genome covered; Fig. 2c). Also, 3.98% and 4.12% of reads from mEnrich-seq are mapped to *N. gonorrhoeae*. Among them, the vast majority (94.66% and 92.26%, respectively) have no GATC sites, representing gDNA molecules that can survive DpnII digestion and PCR amplification; 3.36% and 4.06% of reads have GATC site(s) that overlap with the *N. gonorrhoeae* methylation motif GGTG<u>6mA</u>, thus protected from DpnII digestion; the remaining reads with GATC sites could be due to SNPs or sequencing errors (Fig. 2d; see Methods). Among the reads mapped to the human reference (Fig. 2e), 77.04% and 84.63% have no GATC sites, respectively. Among the 22.96% and 15.37% of reads with at least one GATC sites, the majority (17.33% and 11.99% respectively) of GATC sites are located within genomic regions with simple sequence repeats (see Methods) that tend to form secondary structure, making it less accessible to DpnII digestion^54, 55^. This characterization illustrates that mEnrich-seq not only enriches bacterial taxa of interest, but also carries some by-product reads that are either due to methylation at (partially) overlapping motif sites in background taxa, or depleted restriction motifs in a subset of gDNA molecules. Importantly, the highlight of mEnrich-seq is the enrichment efficiency compared to standard metagenomic sequencing despite by-product reads.

**Fig. 2.**
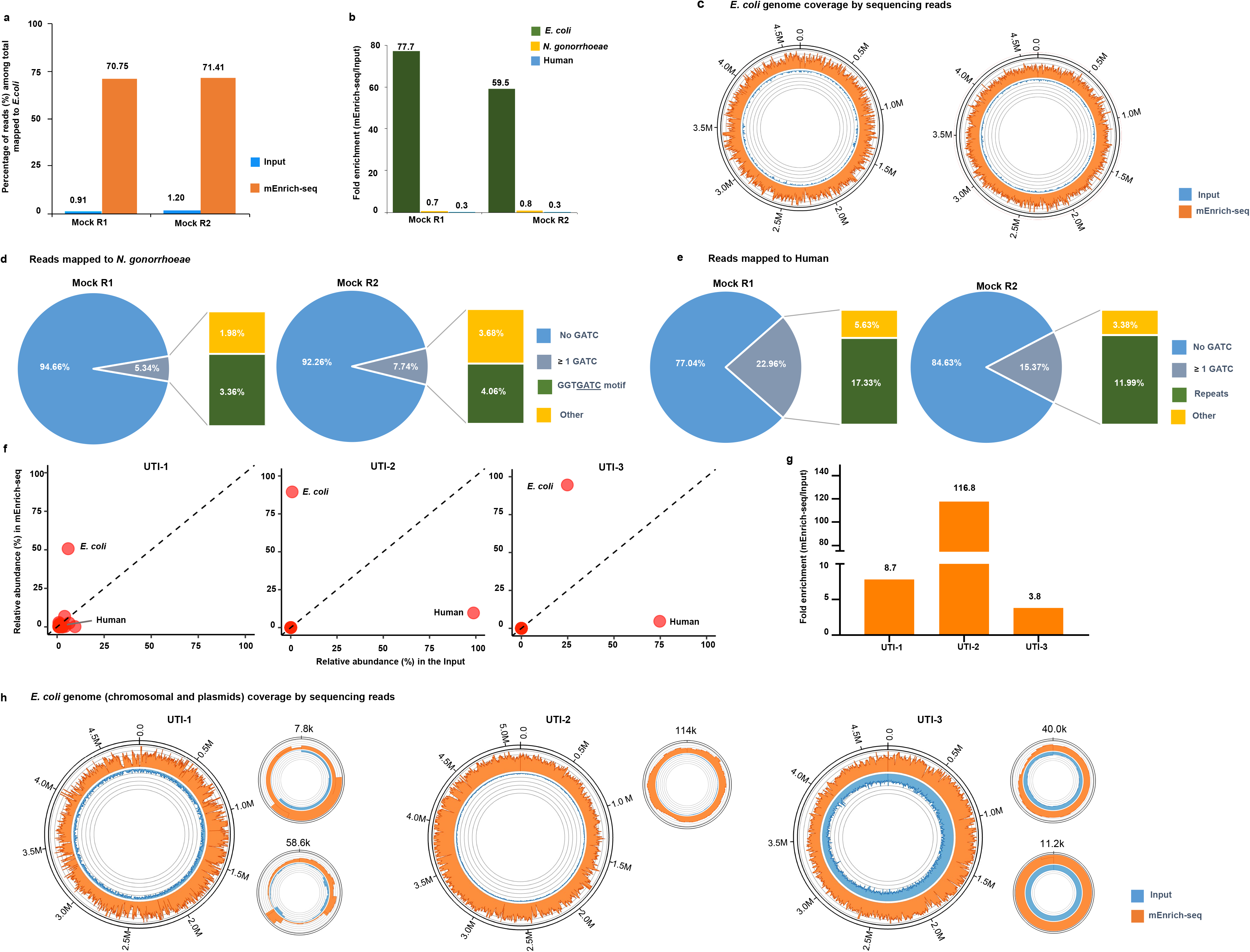
Validation of mEnrich-seq with the mock samples and urine samples. **a**, The percentage of reads mapped to *E. coli* in the two mock replicates; the blue bar represents the input (human 94%, *N. gonorrhoeae* 5%, *E. coli* 1%); the orange bar represents mEnrich-seq. **b,** The fold change (mEnrich-seq vs. input) of percentage reads mapped to *E. coli* (dark green), *N. gonorrhoeae* (yellow) and human genome (blue) in the two mock replicates. **c,** *E. coli* genome coverage by sequencing reads (mEnrich-seq vs. input, with matched total yield). Maximum depth of each circos plot was capped at 95% quantile to ease visualization (Methods). mEnrich-seq reads mapping showed that the *E. coli* genome can be covered without systematic bias recovered for both mock samples (>99.96% genome covered). **d,** Motif frequency on the mEnrich-seq reads that were mapped to *N. gonorrhoeae*. In replicates 1 and 2, 94.66% and 92.26% of reads that mapped to *N. gonorrhoeae* do not have any GATC sites (blue), respectively. The remainder of the reads have ≥ 1 GATC/read (gray), which are further divided into two groups: the reads with GATC sites overlapping with the GGTG<u>6mA</u> motif of *N. gonorrhoeae* (labeled green), and Other (labeled orange), which reflect SNPs or sequencing errors (see Methods). **e,** Motif frequency on the mEnrich-seq reads that mapped to the human genome. In replicates 1 and 2, 77.04% and 84.63% of reads that mapped to human do not have any GATC sites (blue), respectively. The remainder of the reads have ≥ 1 GATC/read (gray), which are further divided into two groups: the reads with GATC sites within repetitive elements that may be inaccessible to RE due to secondary structure (labeled green), and Other (labeled orange), which reflect SNPs or sequencing errors (see Methods). **f,** Scatter plots for three urine samples (UTI-1 to -3) showing the percentage of reads mapped to *E. coli*, human and other taxa from the input (x-axis) vs. mEnrich-seq (y-axis). Across all the three samples, *E. coli* reads are significantly enriched in mEnrich-seq. In UTI-2, and UTI-3, the proportion of reads corresponding to the human genome are significantly decreased due to deletion by mEnrich-seq. **g,** Fold enrichment of *E. coli* reads from mEnrich-seq compared to the input for the three urine samples. A break in the y-axis indicated by a double line indicates a deletion and a change in scale to permit all 3 samples to be graphed in the same plot. **h**, *E. coli* genome (chromosomal and plasmids) coverage by sequencing reads. Reads from mEnrich-seq of urine metagenomic DNA (orange) and the input standard urine metagenomic sequencing (blue) were mapped to the genome of corresponding *E. coli* isolates (sequenced and assembled beforehand for evaluation purpose), respectively. The maximum depth of circos plots were capped at 95% quantile to ease visualization (Methods). mEnrich-seq reads mapping showed that the *E. coli* chromosomes and plasmids can be covered without systematic bias for the three urine samples (> 99.97% of the genome covered).

### Selective enrichment of *E. coli* genomes from urine samples

Building on the efficient enrichment of *E. coli* genome using mock data, we further tested mEnrich-seq on three urine samples (UTI-1 to -3) from patients with urinary tract infection (UTI) (*E. coli* positive urine culture^56^, defined as ≥100,000 CFU [colony forming unit]/ml; Methods). The three UTI samples were sequenced with mEnrich-seq (with DpnII as RE) coupled with Nanopore sequencing. For mEnrich-seq, 1.16-1.58 GB of sequencing data was generated with an average of 233 k reads per sample (mean read length: ∼6 kb, Supplementary Table 2). For comparison with mEnrich-seq, the three urine samples were subjected to standard metagenomic sequencing (labeled as ‘input’), which revealed that UTI-1 contained mostly bacterial DNA, while UTI-2 and UTI-3 were dominated by human DNA (Extended Fig. 1a). We also sequenced the *E. coli* isolates cultured from UTI-1 to -3 and used the *de novo* assembled genomes of these isolates as the background truth for evaluation purposes (see Methods). Using mEnrich-seq, we found a remarkable increase of reads mapped to *E. coli* for all three samples: 8.7-fold for UTI-1 (from 5.88% to 51.14%), 116.8-fold for sample UTI-2 (from 0.77% to 90.00%), and 3.8-fold for UTI-3 (from 24.79% to 93.59%) (Fig. 2f,g). These fold enrichments were further validated using Nanopore Flongle flow cells with lower throughout and cheaper cost to demonstrate flexible utility (Extended Fig.1b,c; Supplementary Table 2). In addition to the efficient fold enrichment, we also evaluated mEnrich-seq in terms of the completeness of *E. coli* genomes recovered from enrichment sequencing based on the comparison with the *de novo* assembled genomes of *E. coli* isolates. mEnrich-seq reads mapping showed that >99.97% of the *E. coli* genomes are covered across all three samples (Fig. 2h).

When analyzing the *E. coli* mEnrich-seq reads, we also observed in the UTI-1 sample substantial nucleotide variation with the genome assembly of the corresponding *E. coli* isolate, suggesting the possibility of multiple *E. coli* strain(s) in this sample. Consistently, we found multiple mEnrich-seq reads with confident mapping to *E. coli* yet harboring genes not included in the genome assembly of the corresponding *E. coli* isolate (Extended Data Fig. 2). To exclude the possibility of low-quality mapped reads from other species, the source of these *E. coli* reads was further confirmed by mapping each read manually to the NCBI-NT database (Methods). This observation is consistent with the increasing recognition that a culture-independent approach may provide additional information that can complement standard urine culture in the pathogen genomes in UTI and other infectious diseases^57–59^ .

For the same cost and sequencing throughput, the efficient enrichment of *E. coli* genomes by mEnrich-seq can facilitate the culture-free study of *E. coli* genomes from urine microbiome with better sensitivity (examining *E. coli* with lower relative abundance) than standard metagenomics. Alternatively, mEnrich-seq can facilitate a more cost-effective examination of urine samples in the study of *E. coli* genomes. In addition to *E. coli*, 6mA at GATC sites are also conserved in the vast majority of Gammaproteobacteria including many enteric pathogens^43, 44, 47, 52, 53^, which makes mEnrich-seq with DpnII applicable to enrich additional enteric pathogens and commensal bacteria as we will describe in the following sections.

### Selectively sequencing *Akkermansia muciniphila* from the infant gut microbiome

*Akkermansia muciniphila*, a gram-negative bacterium colonizing the intestinal tract^60, 61^, has been actively studied for its association with many human diseases, including obesity, type 2 diabetes, and inflammation^62–64^, and also has been linked with patient responses to cancer immunotherapies^65–67^. Because *A. muciniphila* is strictly anaerobic with long doubling time^68^, it is challenging to isolate from fecal samples^69, 70^. RA<u>6mA</u>TTY (R = A or G; Y = C or T), mediated by a 6mA MTase, AmuORF1905P, is one of the 5 methylated motifs in *A. muciniphila* (ATCC-835) according to REBASE (Extended Data Fig. 3a). Using comparative genomics analysis, we found that the MTase is conserved in all the 112 genomes of *A. muciniphila* isolates (Supplementary Table 3). This conserved methylation motif is also validated by *de novo* methylation analysis of two in-house isolates (Extended Data Fig. 3b,c; Methods). This motivated us to apply mEnrich-seq with the RE XapI (or isoschizomer ApoI, digesting non-methylated RAATTY sites), to deplete background DNA and enrich *A. muciniphila* from fecal microbiome without culturing.

**Fig. 3.**
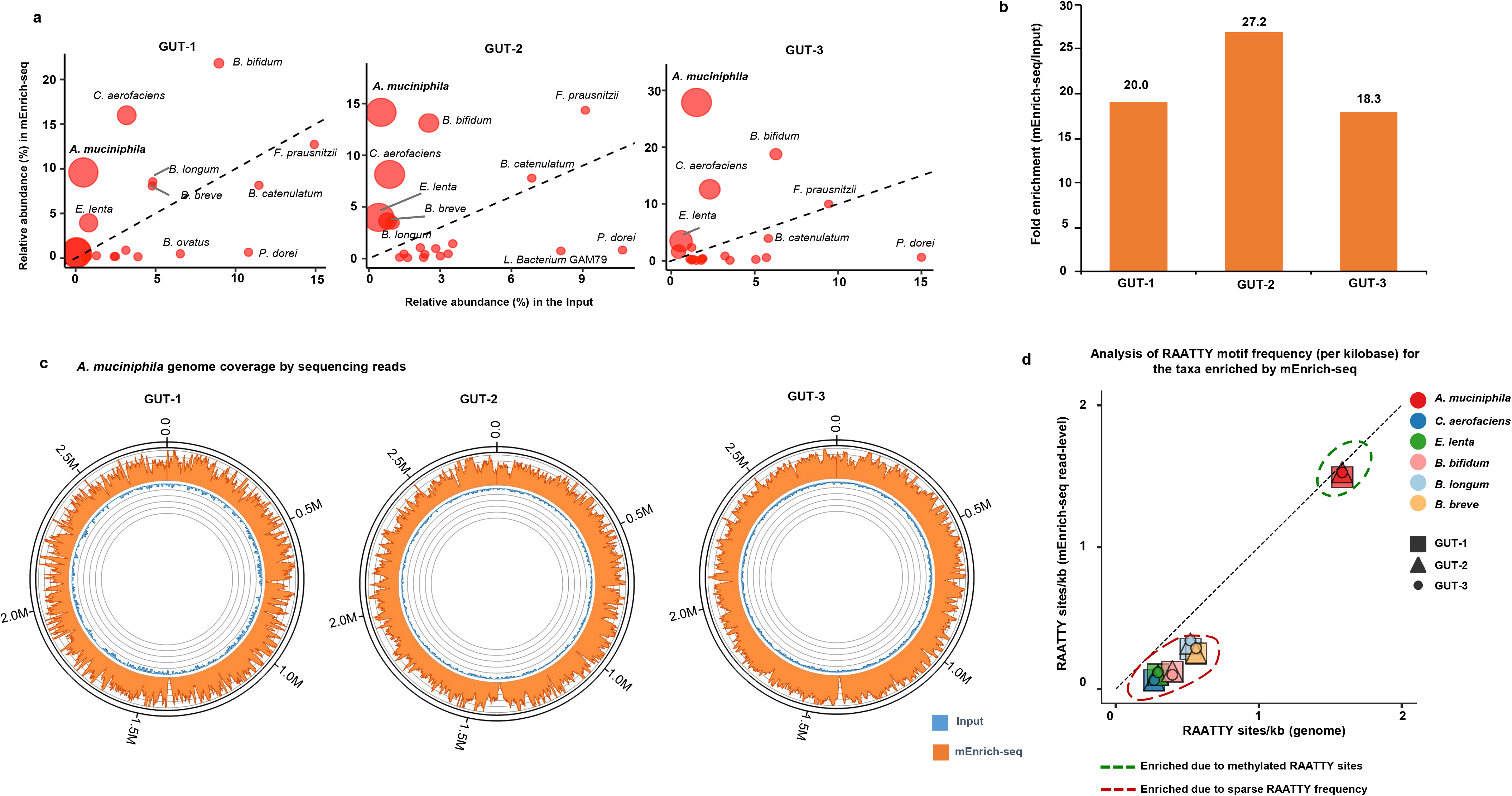
Enrichment of *A. muciniphila* from fecal samples by mEnrich-seq. **a,** Scatter plots for three fecal samples (GUT-1 to -3) showing the percentage of reads mapped to *A. muciniphila* and other taxa from the input (x-axis) and mEnrich-seq (y-axis). To ease visualization, top 50 abundant species, in input or mEnrich-seq, are shown. For the enriched species, the size of the dots represents the fold change by mEnrich-seq over the input. Significant increases in read % are observed for *A. muciniphila* and a few other taxa, going from input to mEnrich-seq samples, while some others are depleted such as *P. dorei*, which undergoes around 10- to 15-fold decrease in read %. **b**, Fold enrichment of reads that map to *A. muciniphila* from mEnrich-seq over the input for GUT-1 to -3. **c,** *A. muciniphila* genome coverage by sequencing reads of GUT-1 to -3. Reads from mEnrich-seq (orange) and the input standard metagenomic sequencing (blue) were mapped to the genome of corresponding *A. muciniphila* isolates (sequenced and assembled beforehand for evaluation purpose), respectively. The maximum depth of circos plot was capped at 95% quantile to ease visualization (Methods). mEnrich-seq reads mapping showed that the *A. muciniphila* genome can be covered without systematic bias for the three fecal samples (>99.7% genome covered). **d,** Analysis of RAATTY motif frequency (per kilobase) for the taxa enriched by mEnrich-seq (XapI as RE) in GUT-1 to -3. X-axis, RAATTY motif frequency on the reference genomes of enriched taxa; Y-axis, read-level RAATTY frequency in mEnrich-seq. Color is used to identify enriched species, while the shapes (square, triangle, or circle) are used to indicate sample of origin. The green dashed line circles *A. muciniphila,* which is enriched due to its methylated RA<u>6mA</u>TTY motif sites. The red dashed line circles the bacterial taxa enriched due to the sparse RAATTY frequency on their genomes.

We applied mEnrich-seq with XapI as the methylation-sensitive RE (XapI cuts non-methylated RAATTY sites, but sensitive to methylated RA<u>6mA</u>TTY sites) to enrich the *A. muciniphila* genome from three infant fecal samples (GUT1-3) with matched isolates for evaluation purpose (see Methods; Supplementary Table 4)^71^. Using standard metagenomic sequencing (labeled as Input), we found that the relative abundance of *A. muciniphila* in GUT-1, GUT-2 and GUT-3 samples were 0.48%, 0.52% and 1.52%, respectively. With mEnrich-seq, we observed a 20.0-fold increase of reads mapped to *A. muciniphila* in GUT-1 (from 0.48% to 9.59%), a 27.2-fold increase in GUT-2 (from 0.52% to 14.16%), and a 18.3-fold increase in GUT-3 (from 1.52% to 27.88%), supporting an efficient and consistent enrichment (Fig. 3a,b). Comparative analysis against the genomes of the *A. muciniphila* isolates from the three infant fecal samples showed that mEnrich-seq was able to cover >99.7% of the *A. muciniphila* genomes (Fig. 3c). Therefore, mEnrich-seq has the advantage of examining *A. muciniphila* genome with 18- to 27-fold (across GUT1-3) efficiency than standard metagenomic sequencing.

In addition to *A. muciniphila*, several taxa showed an elevated proportion of reads compared to the standard metagenomic approach: *Bifidobacterium bifidum*, *Collinsella aerofaciens and Eggerthella lenta etc.* (Fig. 3a). To understand the enrichment of these taxa by mEnrich-seq, we calculated the target motif frequency in the mEnrich-seq data and in the corresponding reference genomes. For *A. muciniphila,* the RAATTY frequency is comparable between the mEnrich-seq reads and RAATTY frequency of the reference genome (Fig. 3d), which is consistent with the enrichment due to methylated RA<u>6mA</u>TTY motif on its genome. In contrast, the frequency of RAATTY in the genomes of *B. bifidum*, *C. aerofaciens* and *E. lenta* is lower, which further decreased among mEnrich-seq reads that mapped to these three species (Fig. 3d). These three genomes with sparse frequency of RAATTY sites were enriched as by-products because of the more efficient depletion of most of the other background genomes with dense yet non-methylated RAATTY sites.

To summarize, mEnrich-seq with XapI as the methylation-sensitive RE (targeting RAATTY) can efficiently enrich *A. muciniphila* (highly conserved RA<u>6mA</u>TTY methylation across all the isolated strains examined), a difficult-to-culture commensal bacterium directly from metagenomic DNA. Since an increasing number of studies highlight the important roles of *A. muciniphila* in human health, diseases, and drug responses^20, 62, 66, 67^, mEnrich-seq can be a useful tool to study the genomes of *A. muciniphila* in a culture-independent, sensitive and cost-effective way, which may facilitate larger-scale analysis of *A. muciniphila* strains and their association with different human diseases.

### Enriching bacterial taxa based on *de novo*discovered methylation motifs

In the previous sections, we have demonstrated the use of mEnrich-seq when the methylation motifs of target bacteria of interest are known *a priori*. Here, we demonstrate the use of mEnrich-seq to enrich gDNA from low-abundance bacteria from complex microbiome based on methylation motifs discovered *de novo* from a microbiome sample of interest.

To facilitate the method evaluation, we built on the adult fecal microbiome sample that has been recently well characterized with comprehensive cultured isolates^42^. In practice, mEnrich-seq does not depend on culture for enriching low-abundant genomes. First, we analyzed a pilot standard metagenomic sequencing of this adult fecal sample (Nanopore sequencing, 7.08 Gb; Methods; Supplementary Table 5). While 36 genomes were assembled (>90% completeness) from high relative abundance taxa using metaFlye (v2.9) ^72^, the others with lower relative abundance could not be well assembled. To rationally enrich less-abundant bacteria with mEnrich-seq, we first needed to know their methylation motifs (Fig. 4a). Using existing tools for *de novo* methylation discovery from microbiome samples^48, 49^ (see Methods), we discovered 112 methylation motifs across 34 strains (from 26 species) with lower abundance (Supplementary Table 6). In this illustration, we prioritized the use of some four 4-mer, 5-mer and 6-mer motifs and REs (excluding GATC, which has been demonstrated in previous sections) across six bacterial taxa with relative low abundance (0.32%-1.62%) for illustration of enrichment by mEnrich-seq.

**Fig. 4.**
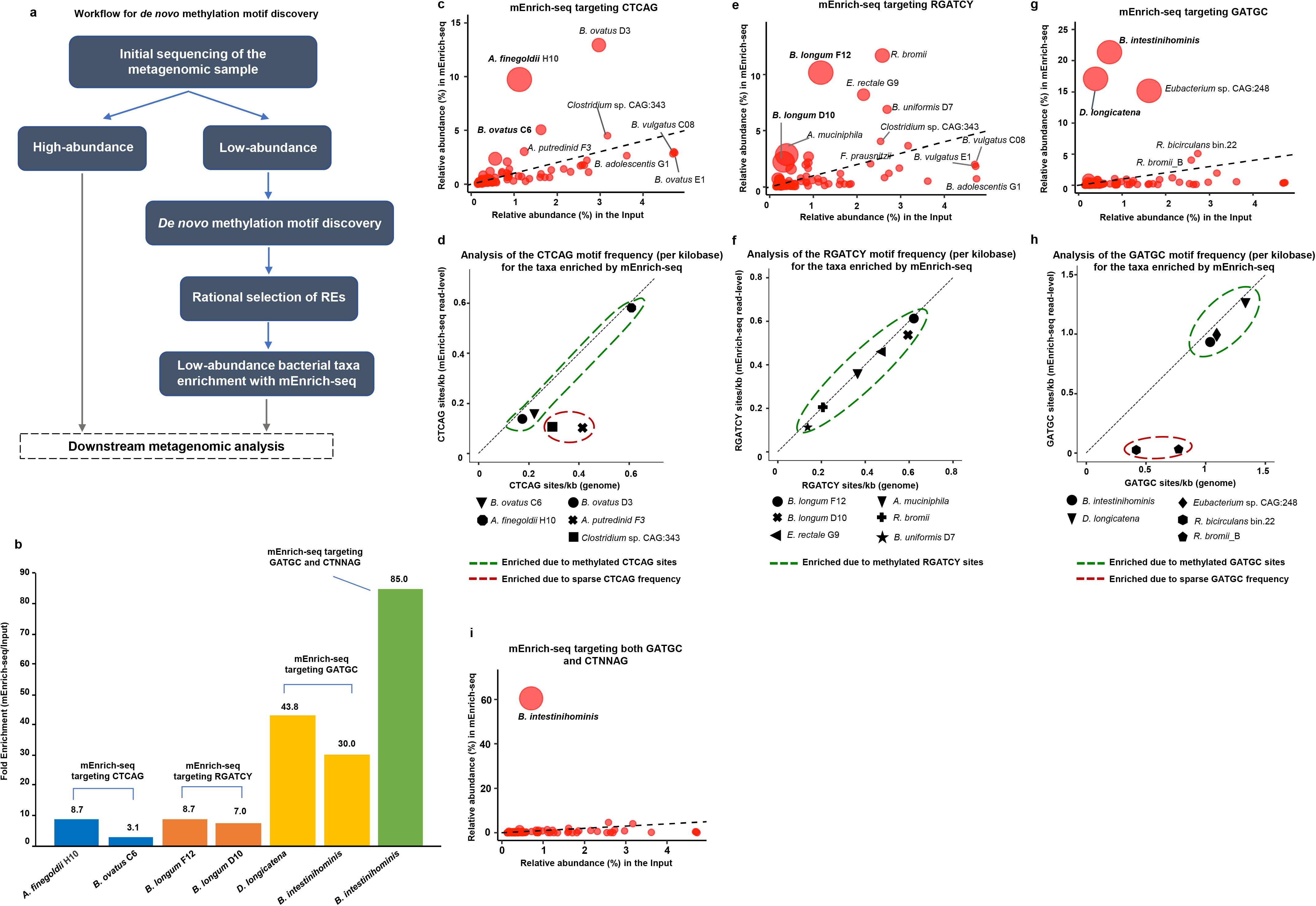
Enrichment sequencing of bacterial taxa based on *de novo* discovered methylation motifs. **a,** Workflow for rational design of mEnrich-seq based on *de novo* motif discovery. An initial metagenomic sequencing was conducted to estimate the relative abundance of different taxa. The goal is to design mEnrich-seq to enrich taxa with relatively low abundance. To do so, we first perform *de novo* methylation motif discovery from the initial metagenomic sequencing data (Methods). Importantly, motif discovery does not need fully assembled genomes, because a partially assembled contig can already inform us about the methylation motifs from a strain. Based on the *de novo* discovered methylation motifs, we selected the REs that rationally deplete the background metagenomic DNA while enriching for bacterial strains with relatively low abundance with mEnrich-seq. **b,** Fold enrichment of six bacterial strains with relatively low abundance; enrichment by mEnrich-seq based on *de novo* discovered methylation motifs. One of the strain *B. intestinihominis,* was further enriched using multiple REs simultaneously to provide greater enrichment (85-fold) than a single RE (30-fold), illustrating the flexible design of mEnrich-seq when multiple methylation motifs are discovered on the strain of interest. **c,** Scatter plot showing the percentage of reads mapped to bacteria from the input (x-axis) and mEnrich-seq using the RE (BspCNI) targeting the CTCAG motif (y-axis). For enriched species, the size of the dots represents the fold change by mEnrich-seq over the input. To ease visualization, top 100 abundant species, in input or mEnrich-seq, are shown. **d,** Analysis of the CTCAG motif frequency (per kilobase) for the taxa enriched by mEnrich-seq (BspCNI as RE). X-axis, CTCAG motif frequency in the reference genomes of enriched taxa; Y-axis, read-level CTCAG frequency in mEnrich-seq. The green dashed line circles the taxa enriched due to their methylated CTC<u>6mA</u>G motif sites. The red dashed line circles the bacterial taxa enriched due to the sparse CTCAG frequency on their genomes. **e,** Similar to **c**, but for the RE targeting the RGATCY motif. **f,** Similar to **d**, but for the RE targeting the RGATCY motif. **g,** Similar to **c**, but for the RE targeting the GATGC motif. **h,** Similar to **d**, but for the RE targeting the GATGC motif. **i,** Similar to **c**, but for the multiple REs targeting the GATGC motif and CTNNAG motif simultaneously.

Two strains (*Alistipes finegoldii* H10 and *Bacteroides ovatus* C6) share the same 5-mer motif CTS<u>6mA</u>G, which can be partially recognized by the methylation-sensitive BspCNI (CTCAG as recognition site) or its isoschizomer BseMII. Aiming to enrich both genomes simultaneously, we applied the mEnrich-seq to the fecal metagenomic gDNA utilizing BspCNI as the RE. As shown in Fig. 4b, encouraging enrichment by mEnrich-seq was achieved for the two genomes: 8.7-fold for *A. finegoldii* H10 (from 1.12% to 9.73%) and 3.1-fold for *B. ovatus* C6 (from 1.62% to 5.05%). In addition to these two genomes, we also found enrichment for two additional genomes: *B. ovatus* D3 (4.3-fold, from 2.97% to 12.94%) and *Alistipes putredinis* F3 (2.4-fold, from 1.23% to 3.01%) (Fig. 4c). Further examination of the CTCAG motif frequency (Fig. 4d) showed that read-level CTCAG frequency in mEnrich-seq data is comparable with CTCAG frequency on the genomes of *B. ovatus* D3 (Fig. 4d), similar to *A. finegoldii* H10 and *B. ovatus* C6 (both with the methylated CTC<u>6mA</u>G sites), reflecting that *B. ovatus* D3 has overlapping methylation at CTCAG sites that resist BspCNI digestion. In contrast, *A. putredinis* F3 was considered as enrichment by-products due to the sparse motifs on their genomes (Fig. 4d).

*Bifidobacterium longum* F12 (input relative abundance 1.17%) and *B. longum* D10 (input relative abundance 0.32%) have the same methylation motif RG<u>6mA</u>TCY (R = A or G; Y = C or T), which can be targeted by the methylation-sensitive RE MflI (or isoschizomers BstX2I, BstYI and PsuI) in mEnrich-seq. Indeed, mEnrich-seq with MflI achieved an 8.7-fold enrichment of the reads mapped to *B. longum* F12 strain (from 1.17% to 10.17%) and a 7-fold enrichment for *B. longum* D10 (from 0.32% to 2.24%) (Fig. 4e). In addition, positive enrichment was observed for *Ruminococcus bromii* (4.5-fold, from 2.58% to 11.69%), *Eubacterium rectale* G9 (3.8-fold, from 2.15% to 8.20%) and *Bacteroides uniformis* D7 (2.6-fold, from 2.69% to 6.89%; Fig. 4e). Further examination of RGATCY motif frequency supported that *R. bromii*, *E. rectale* G9 and *B. uniformis* D7 were enriched owing to methylation at RG<u>6mA</u>TCY sites on their genomes (Fig. 4f), as the read-level RGATCY frequency in mEnrich-seq data are comparable with the RGATCY frequency on their genomes (Fig. 4f).

*Dorea longicatena* and *Barnesiella intestinihominis* share the methylation motif G<u>6mA</u>TGC (reverse strand GC<u>6mA</u>TC). mEnrich-seq with SfaNI as the methylation-sensitive RE provided a 43.8-fold and 30-fold enrichment of reads number in *D. longicatena* (from 0.39% to 17.10%) and *B. intestinihominis* (from 0.71% to 21.34%), respectively. The same data also showed a 9.4-fold enrichment of *Eubacterium* sp. CAG:248 (from 1.62% to 15.17%) (Fig. 4g). Further examination of the target motif frequency showed that *Eubacterium* sp. CAG:248 has a comparable read-level GATGC frequency in mEnrich-seq data as in its genome (Fig. 4h), which supported that *Eubacterium* sp. CAG:248 was enriched similar to *D. longicatena* and *B. intestinihominis* owing to the methylated G<u>6mA</u>TGC motif on their genomes.

Interestingly, in addition to G<u>6mA</u>TGC (reverse strand GC<u>6mA</u>TC), *B. intestinihominis* has one additional methylation motif, CTNN<u>6mA</u>G (N = A, C, G or T), which can be targeted by a mixture of additional REs: AflII (CTTAAG), AcuI (CTGAAG), BpmI (CTGGAG), BpuEI (CTTGAG) and PstI (CTGCAG). This provides us an opportunity to further enhance the enrichment of *B. intestinihominis* with mEnrich-seq. Specifically, we combined the use of SfaNI (targeting GATGC) and the additional five REs targeting the CTNNAG motif into a single mEnrich-seq assay (see Methods), which achieved an 85-fold enrichment of reads mapping to *B. intestinihominis* (from 0.71% to 60.47%), illustrating that mEnrich-seq with combinations of REs targeting multiple motifs can greatly benefit the enrichment efficiency for a specific strain (Fig. 4i).

To summarize, we illustrated that mEnrich-seq can be tailored based on the *de novo* discovered methylation motifs from an adult fecal microbiome sample to enrich bacterial strains with relatively low abundance in the sample. Using three different mEnrich-seq setups (either individual or combinations of REs), we enriched the six rationally targeted genomes. In addition, the same mEnrich-seq setups positively enriched six additional genomes with the same shared methylated motifs (Fig. 4c-g). For the goal of enriching low-abundance genomes from microbiomes, all twelve positive enrichments could have a practical utility to researchers by allowing cost-effectively study specific taxa.

### Broad applicability of mEnrich-seq

To assess the general applicability of mEnrich-seq, we performed a meta-analysis of the methylomes of 4601 bacterial strains (across 1358 species) mapped as of July 18, 2022 (Ref^73^). For each methylome, the list of discovered methylation motifs is maintained at REBASE^73^. On average, there are 3 methylation motifs in each bacterial methylome (Fig. 5a). The number of non-degenerate bases in a motif range from 2 to 9, mostly between 4 and 8 (Fig. 5b). We compared these methylomes with a list of 584 methylation-sensitive REs targeting 209 motifs that are commercially available (Supplementary Table 7) and found that 3130 (68.03%) of the 4601 strains have at least one (26.62% have two or more) methylation motif(s) that can be targeted (enriched by depleting the background metagenomic DNA) by commercially available RE(s). When projected onto a phylogenetic tree (Methods), these 3130 strains have broad taxonomic diversity (Fig. 5c), representing 54.78% of the species examined in this analysis. In this analysis, we focused on REs with recognition motifs that have 4, 5 or 6 non-degenerate bases, as these REs are expected to be readily applicable considering the expected frequency of their target motifs across bacterial genomes: ∼256bp for 4-mers, ∼1kb for 5-mers and ∼4kb for 6-mers. As researchers are actively developing more advanced DNA extraction methods that preserve high-molecular weight gDNA from microbiome samples^74, 75^, we expect the utility of mEnrich-seq to expand to REs with additional target motifs with lower frequency.

**Fig. 5.**
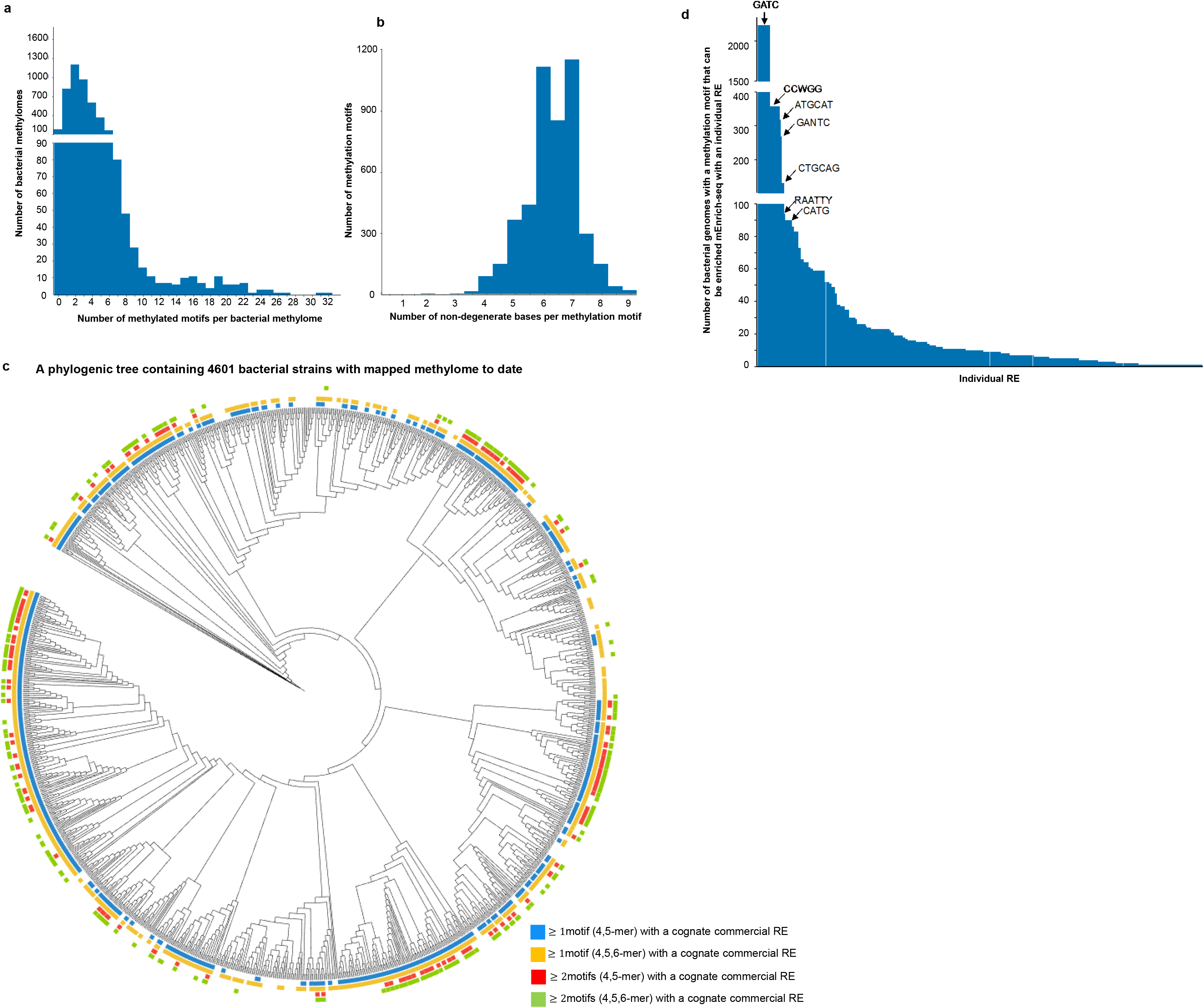
Assessing the broad applicability of mEnrich-seq. **a,** Histogram for the number of methylation motifs in each bacterial genome across the 4601 bacterial methylomes mapped to date (as of 07/18/2022, REBASE). X-axis, number of motifs per bacterium; Y-axis, number of bacterial species having a given number of motifs. **b**, Histogram for the number of non-degenerate bases (excluding ‘N’s) per motif across 4683 motifs. For examples, both GATC and CTNNAG have 4 non-degenerate bases. **c,** A phylogenic tree containing 4,601 bacterial strains with mapped methylomes to date. The bacterial strains that can be targeted (enriched by depleting the background metagenomic DNA) by at least one commercially available REs are highlighted with colors, demonstrating broad distribution across the tree. Number of motifs recognized by commercially available REs (and motif length) for a given bacteria are color coded as detailed in the panel at the bottom right. **d**, Histogram for number of bacteria targeted (enriched by depleting the background metagenomic DNA) by each commercially available RE (please note that a single motif can be targeted by multiple restriction enzymes, namely, isoschizomers). X-axis, each column represents an individual restriction enzyme; Y-axis, number of bacterial genomes per RE target. The most abundant motifs (corresponding to the REs) are indicated by label in the plot.

Some REs have very broad utility as their target motifs are methylated in a large number of bacterial genomes (Fig. 5d). Taking DpnII (methylation-sensitive, targeting GATC) and XapI (and its isoschizomer ApoI, methylation-sensitive, targeting RAATTY) as examples, we have demonstrated in the previous sections their utilities with mEnrich-seq in two specific applications enriching for *E. coli* and *A. muciniphila*, respectively. In addition, mEnrich-seq with DpnII can help enrich additional bacteria in the Gammaproteobacteria class in which G6mATC methylation is highly conserved, while mEnrich-seq with XapI (or its isoschizomer ApoI) can help enrich other bacteria such as *Campylobacter* spp., *Acinetobacter* spp., *Spirochaeta* spp., *Treponema* spp. and *Brachyspira* spp.

## Discussion

In this work, we developed mEnrich-seq, a method that exploits bacterial epigenomes to enrich taxa of interest from metagenomic DNA for selective sequencing. Bacterial DNA methylation provides a natural way for differentiating diverse bacterial taxa from each other. Bacterial DNA methylation has been used for metagenomic binning^48, 49^, but it is only applicable as a post-hoc analysis, after SMRT or Nanopore sequencing. In contrast, mEnrich-seq enabled the use of bacterial DNA methylation to enrich certain taxa of interest *before* sequencing, which provides microbiome researchers with a method to tailor sequencing strategy to best serve their goals. For example, we demonstrated the use of mEnrich-seq to enrich *E. coli* from urine samples and to enrich potentially beneficial species such as *A. muciniphila* from fecal samples. While these two applications built on prior knowledge accumulated from the thousands of bacterial epigenomes mapped to date^45, 73^, we further demonstrated the use of mEnrich-seq along with *de novo* methylation discovery to enrich several low-abundance bacteria from a complex fecal microbiome sample.

mEnrich-seq is broadly applicable and versatile. Based on a meta-analysis of the 4601 bacterial methylomes mapped to date, we estimated that ∼68% of bacterial genomes can be targeted by mEnrich-seq with at least one RE, representing 54.78% of the species examined. In practice, mEnrich-seq may be applicable to a broader diversity of bacterial genomes, because epigenomes mapped to date represent only a small fraction of the bacterial diversity^43, 48^. Consistently, mEnrich-seq enriched several other genomes from urine and fecal samples in addition to the primary species of interest, as they share common methylation motifs being targeted by the methylation-sensitive RE(s) chosen in a specific mEnrich-seq setup. mEnrich-seq is also very versatile in that it can be used along with one or multiple REs (Fig. 1b). Essentially, mEnrich-seq provides a flexible way to dissect a microbiome sample when it is coupled with different REs, the choice of which can be tailored based on taxa of interest in a particular application. In addition, mEnrich-seq is compatible with different sequencing platforms. Although we have mainly used Nanopore sequencing, the core design of mEnrich-seq is also applicable to the PacBio, Illumina and other sequencing platforms. Finally, although we have mainly used mEnrich-seq to study human microbiome samples, this approach is generally applicable to other microbiomes such as environmental samples.

We would also like to note some limitations of mEnrich-seq. First, although mEnrich-seq is broadly applicable and commercially available REs are readily available to all researchers (Supplementary Table 7), future work is needed to expand its use. For example, additional REs can be cloned and purified by individual labs depending on specific needs^76, 77^. In addition, we expect that the increasing use of mEnrich-seq in microbiome applications could also motivate vendors to expand the list of commercially available REs. In the long run, the increasing diversity of REs will be revealed along with the rapidly growing bacterial epigenome studies that characterize both methyltransferases and their cognate REs^43, 45, 48^. Second, not all the reads from mEnrich-seq are from bacterial genomes with the desired DNA methylation. Some by-product reads are expected because some genomes or specific genomic regions can have a very sparse frequency of the RE recognition sites, such as the rare RAATTY motifs on *B. longum* genome due to codon bias (Fig. 3d). Meanwhile, some bacterial reads are enriched because their methylated motifs partially overlapped with the RE recognition motif, as we have characterized in the method evaluations and its applications. The highlight of mEnrich-seq is its high efficacy in the enrichment of taxa of interest over standard metagenomics as we demonstrated across multiple applications. As researchers are actively developing more advanced DNA extraction methods that preserve high-molecular weight gDNA from microbiome samples^74, 75^, we expect the number of undesired reads to be reduced via size selection methods for longer fragments. Finally, mEnrich-seq is applicable to enrich bacteria, archaea and dsDNA phages, but not applicable to ssDNA phages, RNA phages, and parasites, because the latter are not substrates of RM systems or orphan prokaryotic methyltransferases.

mEnrich-seq is complementary to the adaptive sampling method on the Nanopore sequencing platform. Adaptive sampling is very effective in reducing the allocation of sequencing yield to highly abundant host gDNA^30, 78^, or highly abundant bacteria for which enough data have been already collected ^30, 39, 40^. However, with this approach, desired taxa are indirectly enriched by the rejection of reads from highly abundant species, thus the efficiency enrichment by adaptive sampling from latest studies is still modest and insufficient for low-abundant bacteria^30, 78, 79^. In contrast, owing to the high processivity of REs and the diverse methylomes across bacterial taxa, mEnrich-seq has the advantage of more efficiently depleting the vast background gDNA (non-methylated at the targeted RE sites), achieving high fold enrichments of taxa of interest in complex microbiome samples. Adaptive sampling can also select for specific sequences of interest^38–40^ (such as a desired genome); however, it does so by requiring a pre-defined genome sequence hence will unavoidably miss genes unique to a specific strain yet to be determined. In contrast, mEnrich-seq has the advantage of enriching both known and unknown genes in a strain of interest, including mobile genetic elements^48, 49^, because all the genetic contents within a bacterial cell share the same methylation motifs. Therefore, mEnrich-seq and adaptive sampling are complementary.

In summary, mEnrich-seq is a versatile and cost-effective approach for enrichment sequencing of diverse microbial taxa of interest directly from the microbiome. It provides microbiome researchers an additional approach to more effectively study complex microbiomes.

## Acknowledgements

We thank Icahn School of Medicine at Mount Sinai colleagues J. Faith for helping with the characterization of the adult fecal sample and F. Samaroo for helping with urine sample processing. We thank L. Davey and R. Valdivia at Duke University School of Medicine for help with isolation of *A. muciniphila*. We thank M. Blaser Lab at the Rutgers University and other research teams that collected the infant fecal samples for The Early Childhood Antibiotics and the Microbiome (ECAM) study. This work was supported by R35 GM139655 (G.F.) from the National Institutes of Health. G.F. is a Hirschl Research Scholar by Irma T. Hirschl/Monique Weill-Caulier Trust, and a Nash Family Research Scholar. This work was also supported in part through the computational resources and staff expertise provided by the Department of Scientific Computing at the Icahn School of Medicine at Mount Sinai.

## Author contributions

G.F. conceived and supervised the project. L.C. and G.F. designed the methods. L.C. performed all mEnrich-seq experiments and sequencing. Y.K., Y.F., M.N. and A.T. performed the evaluation of mEnrich-seq and analyzed mEnrich-seq data across the applications. M.K. helped with the data analysis of mEnrich-seq of the urine samples. T.K. and M.G. helped with the characterization of urine samples and *E. coli* strains. X-S.Z. helped with the characterization of the infant fecal samples. L.C., M.N., and E.A.M. performed additional sequencing. L.C., Y.F., M.N., X-S.Z. and G.F. performed additional data analyses. L.C. and G.F. wrote the manuscript with inputs and comments from all co-authors.

## Materials and Methods

### Samples

*E. coli* K-12 MG1655 was obtained from American Type Culture Collection (ATCC, cat# 47076, VA, USA) and cultured with Miller Luria Broth (LB) medium, and on LB agar plates (Thermo Fisher Scientific, cat# BP1426-500, and cat# BP1425-500, respectively, MA, USA) at 37°C based on ATCC recommended protocols.

*N. gonorrhoeae* MS11 was obtained from ATCC (cat# BAA-1833) and cultured based on protocols in Dillard, 2011^1^. Briefly, cells were grown in Gonococcal Base medium liquid (GCBL) with Kellogg’s supplements I&II, and on Chocolate II Agar plates (BD, cat# 22126, MD, USA) at 37°C in 5%CO_2_ for 24 h.

Human Lymphoblastoid Cell Line (LCL; Coriell Institute for Medical Research (Coriell), cat# GM 24149, NJ, USA) suspension cell cultures were grown in RPMI-1640 media (GIBCO cat# 11875093, MA, USA) supplemented with 15% Fetal Bovine Serum (GIBCO cat# A3160402) at 37°C in 5% CO_2_. Cells were split at 1:3 to 1:5 ratios when reaching confluency, and were monitored periodically by the MycoAlert Mycoplasma detection assay (Lonza, cat# LT07-118, NJ, USA) for mycoplasma. After 6 passages, cells were harvested for DNA isolation.

Deidentified urinary tract infection (UTI) positive urine samples were obtained after standard urine culture and susceptibility testing performed at the Clinical Microbiology Laboratory (Icahn School of Medicine at Mount Sinai, NY, USA)^2^. No patient identifiable information was collected, only the identified pathogen(s) were provided. Patients provided informed consent to Mount Sinai at collection and further informed consent was not required.

Three infant fecal samples for mEnrich-seq with a RE targeting RAATTY (*A. muciniphila)* were collected from an individual subject from a previous study with Institutional Review Board approval^3^. No personal identifiable information was used in the scope of this work. The matched *A. muciniphila* strains were isolated and cultured by our collaborators from Duke University using the established methods by Becken et al^4^.

The adult fecal sample used for mEnrich-seq with *de novo* discovered methylation motifs was from a previous study with Institutional Review Board approval^5^.

### DNA extraction

DNA from the human LCL cell line GM24149, *E. coli* MG1655, *N. gonorrhoeae* MS11, and urine samples was isolated with the DNeasy Blood and Tissue Kit (Qiagen, cat# 69506, MD, USA) based upon the manufacturer’s instructions, including the optional RNase A treatment provided with the kit. For the human cell line, 2x 10^6^ cells were collected from cell culture by centrifugation for 5 min at 300 x g. The cell pellets were resuspended in 180 μl PBS and 20 μl Proteinase K. After adding 200 μl Buffer AL, the mixture was incubated at 56°C for 30 min. The rest of extraction steps were performed according to manufacturer’s instructions. For *E. coli* MG1655, and *N. gonorrhoeae* MS11, 10^9^ cells were harvested by centrifuging 10 min at 5000 x g. The cell pellets were resuspended in 180 μl Buffer ATL, and then lysed with 200 μl Buffer AL and 20 μl Proteinase K for 30 min at 56°C. The rest of the extraction steps were followed per manufacturer’s instructions.

For urine samples, each 2 ml aliquot of urine was first centrifuged at 5000 xg and 4°C for 30 min, then the pellets were collected in a new tube. Next, the urine pellets were resuspended with 180 μl of the bacterial enzymatic lysis buffer (20 mM Tris pH8.0, 2mM EDTA, 1% SDS and 20 mg/ml lysozyme) and incubated at 37°C for 30 min. Following the enzymatic lysis, 20 μl of Proteinase K and 200 μl Buffer AL was added to the mixture, and it was incubated at 56°C for 30 min. The rest of the extraction steps were followed per manufacturer instructions with one modification, *i.e*., the DNA was eluted with 50 μl AE in the final step to concentrate the DNA.

Fecal DNA was extracted with a QIAamp PowerFecal Pro DNA Kit^6^ (Qiagen, 51804) according to the manufacturer instructions. Briefly, 100 mg of fecal sample, 750 μl of PowerBead Solution and 60 μl of Solution C1 were added to the Bead Tube (provided). After incubation at 65°C for 10 min, the sample was homogenized with the TissueLyser II machine (Qiagen) for 10 min at 25 Hz speed (5 min each side). The homogenized samples were centrifuged to collect the supernatant at 15,000 x *g* for 1 min. The subsequent column-based extraction steps were performed as described in the QIAamp PowerFecal Pro DNA Kit protocol.

#### E. coli isolates from UTI positive samples

The isolates were cultured from the urine samples with confirmed *E. coli* cases^7^. Briefly, 10 μl of urine was spread by disposable inoculating loops (Thermo Fisher Scientific, cat# 22170201) quantitatively onto BD BBL MacConkey agar plates (Becton, Dickinson, and Company (BD), NJ, USA) and incubated for 24 h at 37°C aerobically. For each urine sample, a representative colony of *E. coli* was collected for DNA purification.

### Preparation of the two mock replicates for the initial evaluation of mEnrich-seq

Purified human and bacterial DNA were quantified for concentration with the Qubit dsDNA HS assay (Life Technologies, Q33231, MA, USA), and pooled by adding 94% of human DNA, 5% of *N. gonorrhoeae* DNA and 1% of *E. coli* DNA by mass to create 500 ng of the mock samples. The final ratio of *E. coli, N. gonorrhoeae* and human gDNA was quantified by metagenomic sequencing.

### Detailed methods for mEnrich-seq

Libraries for mEnrich-seq were prepared using the Oxford Nanopore Technology Inc. PCR Barcoding Expansion Pack 1-96 (EXP-PBC096), Ligation Sequencing Kit (SQK-LSK109), and 100-1000 ng of genomic DNA as staring material.

#### Step 1. DNA shearing

Shearing was conducted to establish a uniform size of DNA inserts for better ligation efficiency, to maintain highly consistent molarity estimations for downstream loading into flow cells, and to reduce pore clogging^8^. gDNA was sheared to around 10kb with g-TUBEs (Covaris, cat# 520079, MA, USA) using an Eppendorf 5424 centrifuge at 5000 rpm for 3min at room temperature. The g-TUBE was then inverted and centrifuged for another 3 min to collect the sheared DNA. We concentrated the DNA using 0.5x volume AMPure XP beads (Beckman Coulter, cat# A63881,IN, USA) and eluted with 48 μl nuclease-free (NF) water.

#### Step 2. End Repair and Barcode Adapter Ligation

The fragmented DNA was applied to “End-prep” and “Ligation of Barcode Adapter” steps according to the manufacturer’s instructions. The “End-prep” was performed with 48 μl of sheared DNA in the reaction mixture using NEBNext Ultra II End Repair/dA-Tailing Module and NEBNext FFPE DNA Repair Mix (New England Biolabs (NEB), cat# E7546, M6630, respectively, MA, USA) with incubation at 20°C for 5 min and 65°C for 5 min. After purification with 1x volume of AMPure XP beads, the end-repaired DNA was eluted in 30 μl NF water. Next, the end-repaired DNA was ligated with 20 μl Barcode Adapter (ONT, cat# EXP-PBC096, blue cap) and Blunt/TA Ligase Master Mix (NEB, cat# M0367) for 10min at room temperature, which attached the universal PCR handle to all of the DNA molecules. The Barcode-Adapter-ligated-sample was then purified with 1x volume of AMPure XP beads and eluted in 26 μl NF water. After the “Ligation of Barcode Adapter” step, 20 ng of the PCR Adapter ligated DNA was split into another tube as sample “Input”, while the rest of the sample was used for the restriction enzyme digestion step of mEnrich-seq.

#### Step 3. Restriction enzyme (RE) digestion

RE digestion for mEnrich-seq was conducted following the manufacturer’s instructions. For DpnII digestion, 10 U of enzyme (NEB, cat# R0543) was used in a 50 μl-reaction containing 5 μl of NEBuffer r3.1 (10X, NEB, cat# B7203), 26 μl of adapter-ligated DNA and NF water to 50 μl. The reaction system was incubated at 37°C for 30 min. For BspCNI (NEB, cat# R0624S), 10 U of the enzyme was used in the 50 μl-reaction containing 5 μl of rCutSmart Buffer (10X, NEB # B6004S) for 30 min-incubation at 37°C. 10 U of XapI (Fisher Scientific, ER1381) was added in 50 μl-reaction system supplied with 5 μl Buffer Tango (10X, Fisher Scientific, BY5) and incubated for 30 min at 37°C. When using SfaNI (NEB, cat# R0172S) for digestion, 10 U of enzyme was used in 50 μl reaction system with 5 μl of NEBuffer r3.1, and incubated for 30 min at 37°C. MflI (Takara, cat # 1070A) was used with 10 U in the 50 μl-reaction system containing buffer L, and incubated at 37°C for 30 min.

When using multiple enzymes, it’s critical to first check whether the enzymes work in the same optimal buffer and temperature as recommended by the manufacturer. In our test, we used 6 enzymes in combination, SfaNI, AflII (NEB, cat# R0520S), AcuI (NEB, cat# R0641S), BpmI (NEB, cat# R0565S), BpueI (NEB, cat# R0633S) and PstI-HF (NEB, cat# R3140S). Among them, only SfaNI works in NEBuffer r3.1 at 100% efficiency. To make sure all enzymes work optimally, the DNA was firstly digested with 20 U SfaNI in NEBuffer r3.1 for 30 min at 37°C, then the digestion system was purified with 1.8X AMpure beads. The eluted DNA (20 μl) was further digested for 30 min at 37°C with a 5 μl-mixture of AflII, AcuI, BpmI, BpueI and PstI-HF (1 μl of each).

#### Step 4. Gel purification

The RE-digested products were loaded into a 1.5% agarose gel (Thermo Scientific, cat# R0492, MA, USA) pre-stained with SYBR Safe DNA gel stain (Invitrogen, cat# S33102, MA USA) for gel-based size selection. DNA ladder (NEB, cat#N3238S) was used as a comparison for size estimation and each gel was visualized under UV light on a ChemiDoc XRS+ System (Bio-Rad, CA, USA). The DNA band at 5 to 10kb size was collected for extraction with a NucleoSpin Gel and PCR Clean-up purification kit following the manufacturer’s instructions (Macherey-Nagel, cat# 740609.250, Germany), and was eluted with 15 μl NF water.

#### Step 5. PCR amplification

To add the barcode to each sample, the gel-purified DNA was amplified with the barcoded primers from the barcoding expansion pack mentioned above (EXP-PBC096, white caps, ONT). For PrimeSTAR GXL Premix (2x, Takara Bio, cat# R051A, Japan), a 50 μl -reaction system was prepared as follows: 25 μl of Premix, 1 μl of barcoded primer (10uM, white tube),15 μl purified DNA and 9 μl water. The PCR reactions were pre-heated at 98°C for 1 min followed by 15-18 cycles of 98°C for 15 s, annealing 62°C for 15 s and extension at 68°C for 8 min, with a final extension at 68°C for 10 min on the ABI Veriti thermal cycler. The PCR products were purified with 0.5x volume of AMPure XP beads. Before moving to library prep, the purified PCR products were checked with an agarose gel for a distinct band at 5-10 kb in size (Extended Fig. 4-7).

#### Step 6. Library prep and sequencing

The purified barcoded PCR products were pooled together for “DNA repair” and “end-prep” step. Briefly, 47 μl of pooled PCR products were incubated at 20°C for 15min and 65°C for 15min, in a 60-reaction system containing 3.5 μl of NEBNext FFPE DNA Repair Buffer, 2 μl of NEBNext FFPE DNA Repair Mix, 3.5 μl of Ultra II End-prep reaction buffer and 3 μl of Ultra II End-prep enzyme mix. The purified DNA was ligated to Adapter Mix (ONT, SQK-LSK109) in the presence of Ligation Buffer (ONT, SQK-LSK109) and NEBNext Quick T4 Ligase (NEB, cat# E6056) for 30 min at room temperature. After beads purification, the adapter ligated DNA was ready for sequencing on ONT flow cells.

### Compatibility between mEnrich-seq and other sequencing platforms

Although we focused on the use of Nanopore sequencing in this manuscript, mEnrich-seq is compatible with other sequencing platforms, because the barcoded PCR products from ‘step 5’ in the detailed methods are essentially double-stranded DNA. For PacBio platform users, after purification with AMPure PB beads (Pacific Biosciences, #100-265-900), the PCR products from “Step 5” are ready for library preparation following the “Preparing multiplexed amplicon libraries using SMRTbell prep kit 3.0” (PN 102-359-000 REV 02 SEP2022). It’s recommended to use the barcode kit by PacBio to achieve the best sequencing result. The reads from mEnrich-seq are also computational applicable for data analysis by trimming the barcode adapter sequences (see “Guidance for broad applications of mEnrich-seq”).

For the users of the Illumina platforms, as Illumina is a short-read platform, the barcoded PCR products from ‘**Step 5**’ will firstly be purified and adjusted to shorter insert size (*e.g.,* by sonication). Then, the fragmented DNA molecules are ready for users to prepare the library with suitable kits compatible with various Illumina platforms. Compatible barcoding kits are required for a good sequencing result. The Barcode adapter sequence from the mEnrich-seq reads can be trimmed using computational methods for further analysis.

#### Guidance for broad applications of mEnrich-seq

##### Barcode adapter

As the Barcode adapter ligation step is prior to the RE digestion during the library preparation, the DNA from targeted taxa will not be amplified by PCR if the barcode adapter is cut during the RE digestion step. Therefore, it’s critical to avoid a chosen RE cutting the sequence of the Barcode adapter or the junction between the adapter and metagenomic DNA. We suggest to check the underlined sequence of the Barcode adapter (reverse complementary to each other) for any possible RE motif site. If there’s a recognition sequence for the chosen RE, the bases can be substituted to avoid the cut. The sequences are as follows:

top sequence, 5’-TTTCTGTTGGTGCTGATATTGC<u>GGCGTCTGCTTGGGTGTTTAACC</u>*T-3’ and bottom sequence 5’-<u>GGTTAAACACCCAAGCAGACGCC</u>GAAGATAGAGCGACAGGCAAGTA-3’. The top and bottom sequence are added to the annealing buffer (10 mM Tris, pH 7.5, 50 mM NaCl and 1 mM EDTA) to a concentration of 10 μM and annealed with the program: 95 °C for 2 min, −0.1 C/sec for each of 800 s (∼13 min) and hold at 4°C. The barcoded adapter is now ready to use and compatible with the original protocol. **Selection of RE.** The RE can be found through the Enzyme Finder (https://enzymefinder.neb.com/#!#nebheader) based on the chosen target motifs. For the chosen RE, it’s important to check its methylation sensitivity result provided by the manufacturer in REBASE (http://rebase.neb.com/rebase/rebase.html), which is commonly tested with modified oligos, and even better to re-confirm the sensitivity upon purchase.

### Library preparation for the standard metagenomic sequencing (Input)

For the mock, urine and the infant fecal samples, libraries of the standard metagenomic sequencing (Input) for comparison with mEnrich-seq were prepared with the 20 ng DNA split from Barcode Adapter ligation step (at the end of ‘Step 2’ in mEnrich-seq). The Input sample was first amplified with barcoded primer. The PCR master mix consisted of 1 μl PCR barcode (10 μM, ONT), 25 μl PrimeSTAR GXL Premix (2x, Takara Bio, cat# R051A, Japan), 20 ng DNA and NF water to a total volume of 50 μl. The PCR programs were as follows: pre-incubation at 95°C for 3 min, amplification for 14 cycles at 95°C for 15 s, 62°C for 15 s, and extension at 65°C for 10 min, with a final extension at 65°C for 10 min. The PCR products were checked with an agarose gel for successful amplification (bands at ∼5 to 10 kb, Extended Fig. 4-7). After 0.5x volume bead purification, the purified barcoded PCR products were used for “repair and end-prep” step in the ONT protocol. Following the “Adapter ligation and clean-up” step, the library was quantified by Qubit and ready for sequencing.

### Library preparation for other microbiome samples and isolates

The data of adult fecal microbiome sample were generated from another study. Briefly, 1000 ng of the DNA was used for library preparation using SQK-LSK109 kit from ONT, following the Native Barcoding genomic DNA protocol (NBE_9065_v109_revZ_14Aug2019). Briefly, the purified DNA was sheared with g-TUBE, and then was added to “End-prep and FFPE repair” reaction system and incubated at 20°C for 5 min and 65°C for 5 min. Following the DNA purification step, the eluted DNA was added to the sequencing “Adapter ligation” reaction system, containing Ligation Buffer (LNB), NEBNext Quick T4 DNA Ligase and Adapter Mix. After purification using 0.5x beads, the prepared library was quantified with Qubit, and 40 fmol of the library was loaded onto a MinION cell by manufacturer’s instructions.

For the *E. coli* isolates from urine samples, we followed the library preparation protocol from Oxford Nanopore Technologies plc (ONT), “Native barcoding genomic DNA” (with EXP-NBD104, EXP-NBD114, and SQK-LSK109, ONT, NY, USA) with one modification for high molecular weight shearing. Briefly, 5 μg purified gDNA of *E. coli* isolates was diluted in 250 μl EB buffer (10 mM Tris-Cl, pH8.0), then the DNA was sheared with a 29 G needle for 20 passes^9^. This led to DNA samples with an N50 of approximately 20 kb. The sheared DNA was purified with 0.5x volume of AMPure PB beads (Pacific Biosciences of California, Inc.), and then was eluted with 51 μl nuclease-free water. After quantification by Qubit, 1 μg of sheared DNA was used for further library preparation.

The sequencing data of *A. muciniphila* isolates from the infant fecal samples were collected from another study. In brief, the SMRTbell libraries of *A. muciniphila* strains were prepared according to the manufacturer’s instructions, and then sequenced on the Sequel instrument. 6mA methylation motif discovery from *A. muciniphila* isolates were performed with the build-in pipeline pb_basemods from *SMRTlink v10.0*.

### Nanopore sequencing and base calling

Sequencing was primarily performed with both MinION flow cell R9.4.1 and Flongle flow cell R9.4.1 (both from ONT), and ONT MinKNOW software (v21) was used for sequencing raw data collection. For the MinION flow cell, 50 fmol of library was loaded onto flow cell according to manufacturer’s instructions and the MinION was run for 72 h to collect the raw data. For the Flongle flow cell, 20 fmol of library was loaded and run for up to 24 h to collect the raw data. All Nanopore sequencing reads were base called using guppy (v5.0.7) with the SUP model dna_r9.4.1_450bps_sup.cfg.

### Evaluation of mEnrich-seq of the two mock replicates

#### Fold enrichment and completeness of *E. coli*

The genome references of *E. coli* K12 MG1655 (RefSeq accession CP014225.1), *N. gonorrhoeae* MS11 (RefSeq accession CP003909.1), and human (GRCh38) were combined as the mock reference. Repeat annotation of human genome was obtained from RepeatMasker database (https://www.repeatmasker.org/genomes/hg38/RepeatMasker-rm405-db20140131/hg38.fa.out.gz). The fold enrichment of *E. coli* was calculated based on *E. coli* reads from mEnrich-seq and Input samples that were mapped to the mock reference using *minimap2* (v2.24-r1122) with *-x map-ont*. For genome completeness, the coverage of the *E. coli* was calculated using read depths across 100bp bins across the *E. coli* genome. For Circos plots, Nanopore reads from paired Input and mEnrich-seq samples were normalized to the same yield by random subsampling using *seqtk* (v1.2-r94) with 2-pass mode. Reads after normalization were mapped to the mock reference using *minimap2* (v2.24-r1122) with *-x map-ont*. Sliding windows with intervals of 2.5kb and step sizes of 1.25kb were defined across the *E. coli* genome using *bedtools* (v2.29.2) with *-w 2500 -s 1250*. Reads mapped to defined windows were counted using *bedtools multicov*. Circos plots were drawn using *circos* (v0.69-6). To ease visualization, maximum depth of each circos plot was capped at the 95% depth quantile.

#### Motif analyses in *N. gonorrhoeae* and human

Nanopore reads from the Input and mEnrich-seq samples were mapped to the mock reference using *minimap2* (v2.24-r1122) with *-x map-ont*. Reads of *N. gonorrhoeae* and human mEnrich-seq were extracted using *samtools* (v1.15.1). To reduce the possibility of low-quality mis-mapped reads from other species, non-primary and supplementary alignments were not included for the motif analysis using *samtools* (v1.15.1) with *view -F 2308*. Considering the moderate Nanopore sequencing error rate, analysis of motif (G<u>A</u>TC and GGTG<u>A</u>TC) frequencies and overlapping of G<u>A</u>TC motif and human genomic repeats were counted based on mapped reference segment of a read instead of the read itself.

### Evaluation of mEnrich-seq of the urine samples

#### Fold enrichment and completeness of *E. coli*

The references of *E. coli* isolates were assembled using Flye (v2.8.1-b1684) with *--nano-raw --genome-size 5m*. *E. coli* reads were extracted by mapping the Input and mEnrich-seq Nanopore reads to the references of the *E. coli* isolates using *minimap2* (v2.24-r1122) with *-x map-ont*. Non-*E. coli* reads were then classified using Kraken2 (v2.0.8-beta) with the k2_pluspf_20200919 database. For genome completeness, the coverage of the *E. coli* was calculated using read depths across 100bp bins across the *E. coli* genome assembled from the isolates. For Circos plots, Nanopore reads from paired Input and mEnrich-seq samples were normalized to the same yield by random subsampling using *seqtk* (v1.2-r94) with 2-pass mode. Reads after normalization were mapped to the references of the *E. coli* isolates using *minimap2* (v2.24-r1122) with *-x map-ont*. To draw the circos plots, sliding windows with intervals of 2.5kb and step sizes of 1.25kb were defined across the *E. coli* references using *bedtools* (v2.29.2) with *-w 2500 -s 1250*. Reads mapped to defined windows were counted using *bedtools multicov*. To exclude low-quality mapped reads from redundant genomic locations or other species, *e.g.*, high similar reads between multiple copies of rRNA, read depths were capped at the 95% depth quantile for each chromosome (*i.e.* the longest circled contig 4∼5Mb) and plasmid (*i.e.* short circularized contigs) for Circos plots drawing using *circos* (v0.69-6).

### Evaluation of mEnrich-seq of the infant fecal samples

#### Fold enrichment and completeness of *A. muciniphila*

Reads from the Input and mEnrich-seq were assessed using *Kraken2* (v2.0.8-beta) classification with the k2_pluspf_20200919 database. To check the completeness of the *A. muciniphila,* Nanopore reads were mapped to the *A. muciniphila* isolate reference and calculated with reads depth in 100bp bins across the genome. Before Circos plots, Nanopore reads from paired Input and mEnrich-seq samples were normalized to the same yields by random subsampling using *seqtk* (v1.2-r94) with 2-pass mode. Reads after normalization were mapped to the reference of *A. muciniphila* using minimap2 (v2.24-r1122) with *-x map-ont*. To draw the circos plots, sliding windows with intervals of 2.5kb and step sizes of 1.25kb were defined across the *A. muciniphila* isolate reference using *bedtools* (v2.29.2) with *-w 2500 -s 1250*. Reads mapped to defined windows were counted using *bedtools multicov*. To exclude the low-quality mapped reads from redundant genomic locations, *e.g.,* high similar reads between multiple copies of rRNA, read depths were capped at the 95% depth quantile of the whole genome for Circos plots drawing using *circos* (v0.69-6).

#### RAATTY motif analysis

For the species used for motif analysis, all genome references except *A. muciniphila* were retrieved from NCBI, including *C. aerofaciens* (RefSeq genome sequence GCF_002736145.1), *E. lenta* (GCF_021378605.1), *B. bifidum* (GCF_000273525.1), *B. longum* (GCF_000196555.1), *B. breve* (GCF_001281425.1). The matched *A. muciniphila* reference was obtained from the isolated strain as previously described. Nanopore reads from the mEnrich-seq samples were mapped to the reference of each species using *minimap2* (v2.24-r1122) with *-x map-ont*. To reduce the possibility of low-quality mis-mapped reads from other species, non-primary and supplementary alignments were not included in motif analysis using *samtools* (v1.15.1) with *view -F 2308*. Considering the moderate sequencing error rate of Nanopore sequencing, analysis of RAATTY motif frequencies in mEnrich-seq were counted based on mapped reference segment of a read instead of the read itself. The genome frequency for RAATTY motif on each species was counted with custom PERL script.

### *De novo* motif discovery from the adult fecal microbiome

*De novo* methylation motif discovery from the fecal microbiome DNA sample was based on PacBio sequencing data (SEQUEL II). Briefly, the DNA SMRTbell library was prepared according to the manufacturer’s instructions, then the library was annealed to sequencing primer together with the Sequel 2.1 polymerase, followed by loading to the 8M sequencing chip at concentration of 125-175pM. 2.5 Gb CCS data were generated with 400k ccs reads and 6,341bp as median read length.

The raw SMRT reads pre-processing was performed with *SMRTlink v8.0* (Pacific Biosciences). Raw multiplexed files were first de-multiplexed with lima. Circular Consensus reads (CCS reads) were generated using *CCS* software (v4.0.0) with *--min-rq 0.99* based on subreads. CCS reads were assembled using *metaFlye* (v2.9) with *--pacbio-hifi*. The assembly was then binned and refined with the *metaWRAP Binning* and *Bin_refinement* module^10^, which could combine the results from three metagenomic binning software - *MaxBin2*, *metaBAT2*, and *CONCOCT*. We measured the completeness and the contamination level using *CheckM* (v.1.1.3). The metagenomic profile computing and graphics were performed by R (v.4.2.1). For evaluation purpose, the genome assemblies from bacteria isolates from a previous study^5^ along with *de novo* assembled metagenomic contigs were used as the reference database for the following nanopore reads assignments and motif analyses.

The *de novo* methylation motif discovery was performed following the previous work^11^. Briefly, the bins of species/strain level with low completeness (10% ∼ 80%) were selected for further analysis. We mapped the SMRT subreads to metagenomes using *pbalign* (v0.4.1) with default parameters and separated the alignments into each species/strain based on the binning results. IPD ratio were calculated with *ipdSummary* (v2.4.1) from the alignments for each species/strain. The *de novo* motifs were detected by using *motifMaker* (in *SMRTlink v8.0* package) with default parameters. Only the motifs with detection percentage > 60% were kept for the following analysis (Supplementary Table 2).

### Evaluation of mEnrich-seq for the enrichment of low-abundance species from the adult fecal sample

#### Reads assignments

Nanopore reads were first mapped to the reference database described above using *minimap2* (v2.24-r1122) with *-x map-ont*. To reduce the possibility of low-quality mis-mapped reads from other species, non-primary and supplementary alignments were not included for reads assignment using *samtools* (v1.15.1) with *view - F 2308*. Other reads were then classified using *Kraken2* (v2.0.8-beta) with the k2_pluspf_20200919 database.

#### Motif analysis

For each targeted genome, Nanopore reads from the mEnrich-seq samples were extracted from the above read assignment results. Considering the moderate sequencing error rate of Nanopore sequencing, analysis of motif frequency in mEnrich-seq were counted based on mapped reference segment of a read instead of the read itself. The genome frequency for a motif on specific species were counted with custom PERL script.

### Assessing the broad applicability of mEnrich-seq

The REBASE database is constantly updated with mapped bacterial methylomes to date. For each bacterial methylome, a list of methylation motifs was reported (http://rebase.neb.com/rebase/rebase.html) ^12^. We extracted all 4-mer to 6-mer methylation motifs and their corresponding restriction endonucleases (REs) from 4,644 bacterial methylomes (as of 7/18/2022). Among 4,644 PacBio organisms, we selected the 1,358 most complete genomes as representatives when multiple strains from the same species are present. The protein sequences from the 1,358 species level genomes were extracted to reconstruct a phylogenetic tree using *PhyloPhlAn* (v3.0)^13^. Visualization and annotation of the phylogenetic tree was performed by *ITOL* (https://itol.embl.de/).

