## Supplementary Figures for "mEnrich-seq: Methylation-guided enrichment sequencing of bacterial taxa of interest from microbiome"

### Extended Figures

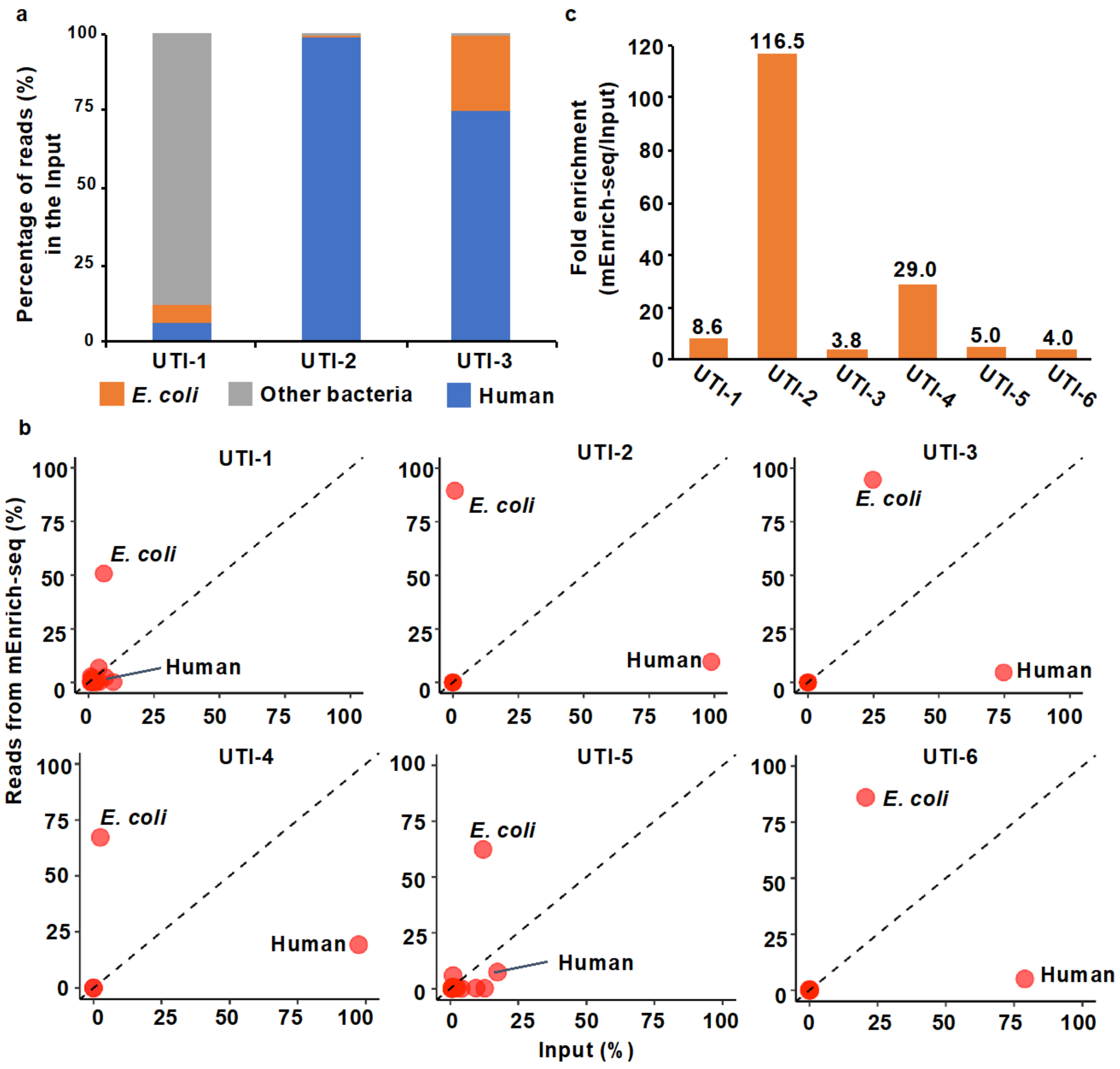


**Extended Fig. 1 mEnrich-seq of urine samples using Nanopore Flongle flow cells. a,** Three urine samples were sequenced as the Input: UTI-1 contained mostly bacterial DNA, while UTI-2 and UTI-3 were dominated by human DNA. **b,** Scatter plots for six urine samples (UTI-1 to -6) showing the percentage of reads mapped to *E. coli*, human and other taxa from the input (x-axis) vs. mEnrich-seq (y-axis)**.** The UTI-1 to -3 samples demonstrate consistent patterns as shown in Fig. 2 in the main text (MinION flowcell); the additional three UTI urine samples (UTI -4 to -6) are included to illustrate the consistent enrichment of *E. coli* by mEnrich-seq. Across all six samples, *E. coli* reads are significantly enriched in mEnrich-seq. In UTI-2, UTI-3, UTI-4, and UTI-6, the proportion of reads corresponding to the human genome are significantly lower due to cleavage by DpnII in mEnrich-seq. **c,** Fold enrichment of *E. coli* reads from mEnrich-seq compared to the input for the six urine samples.

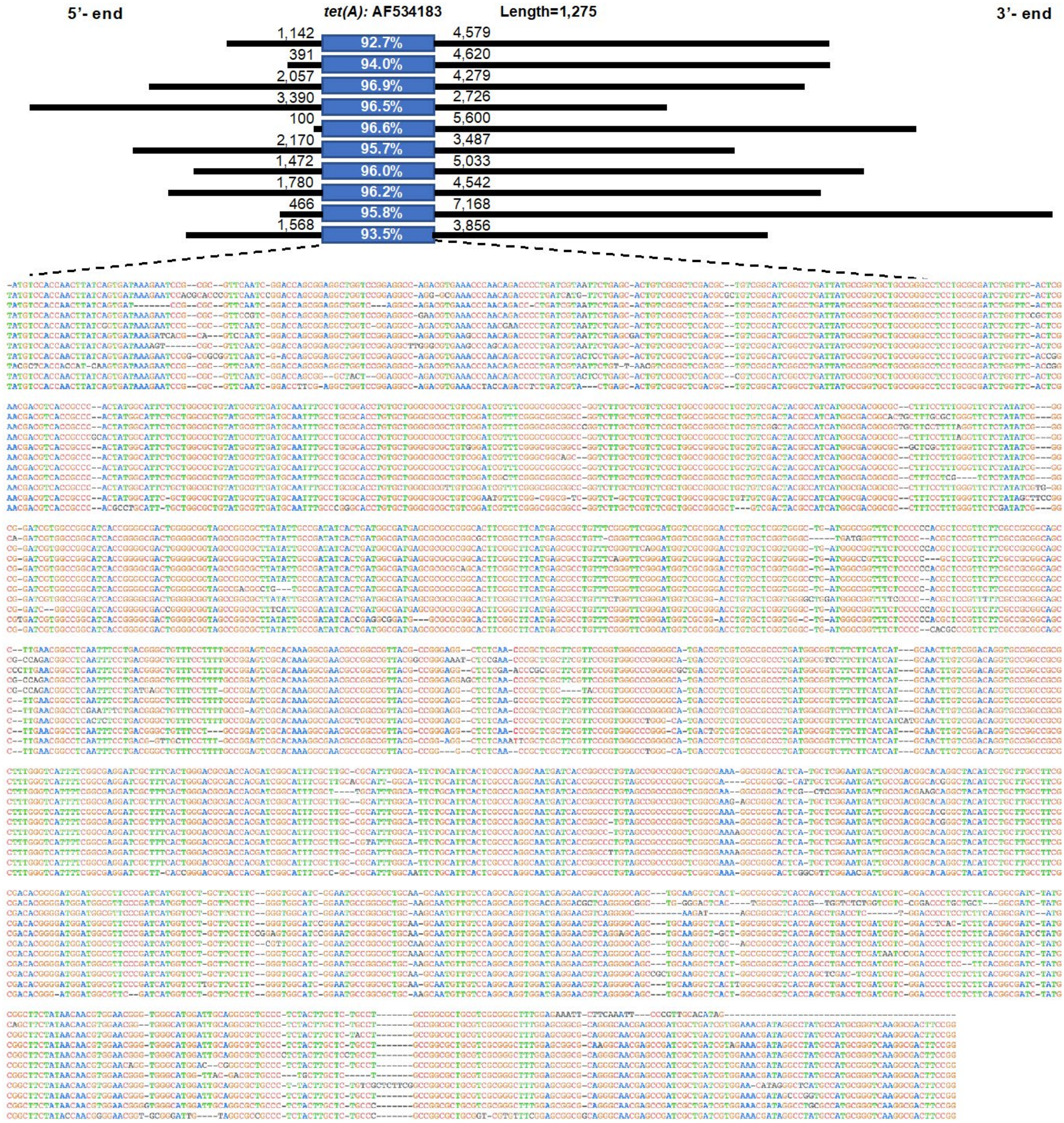


**Extended Fig. 2 mEnrich-seq data of UTI-1 include reads that have high-confidence mapping to *E. coli* yet carry additional genes beyond the isolated strain from standard urine culture.** Top: 10 reads with highly confident mapping to *E. coli* from mEnrich-seq of UTI-1 urine sample, consistently supporting a *tet(A)* resistant gene (AF534183, blue boxes). For each read, lengths of the sequences flanking the resistance gene were shown on both sides. The identities between the reads and AF534183 were shown inside the blue boxes. Bottom, reads alignment to AF534183 was performed with Clustalw2 and colored with MView.

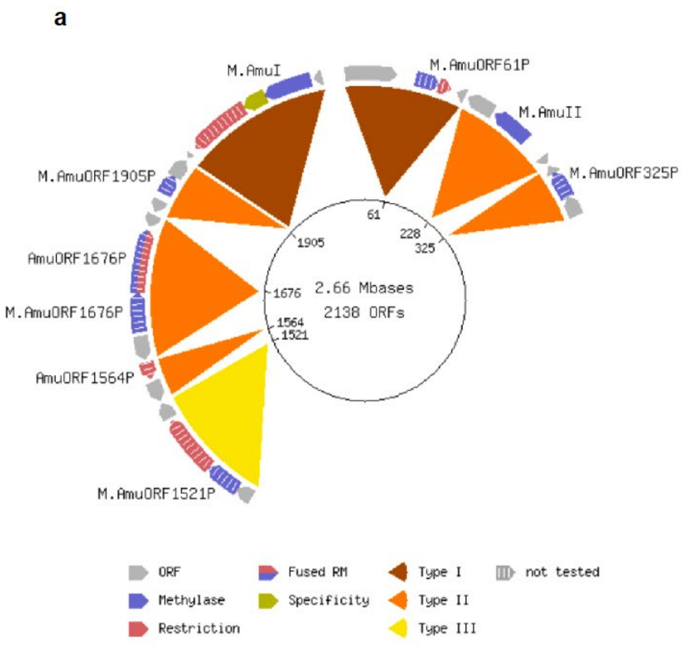


| Motifs | Enzymes |
|---|---|
| RAATTY | M.AmuORF1905P |
| CAAGAA | M.AmuII |
| GTRACC | |
| TCMKGA | |
| AACNNNNNNTAC | AmuI |

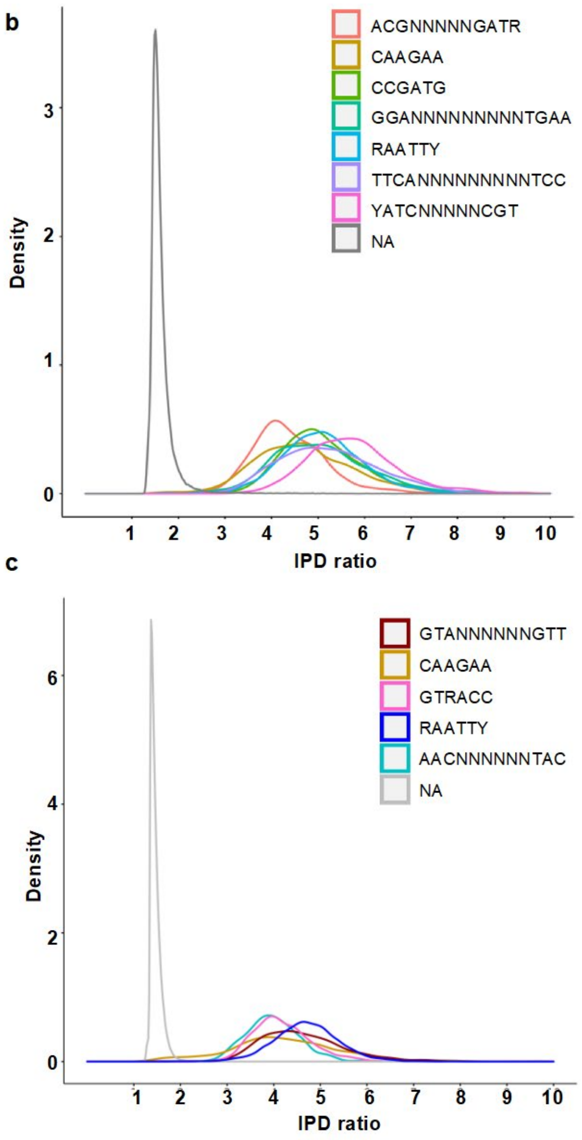


**Extended Fig. 3 Confirmation of DNA 6mA methylation at RA6mATTY motif sites using PacBio sequencing of two *A. muciniphila* strains. a,** RA6mATTY (mediated by MTase AmuORF1905P) is one of the five methylated motifs detected in *A. muciniphila* (ATCC-835) by PacBio sequencing as summarized in REBASE. **b,** DNA 6mA methylation at RAATTY motif sites (light blue) is confirmed based on the high IPD ratios observed in PacBio sequencing data of the *A. muciniphila* strain isolated from the infant fecal sample GUT-3. **c,** DNA 6mA methylation at RAATTY motif sites (purple) is confirmed based on the high IPD ratio observed in PacBio sequencing data of another *A. muciniphila* strain.

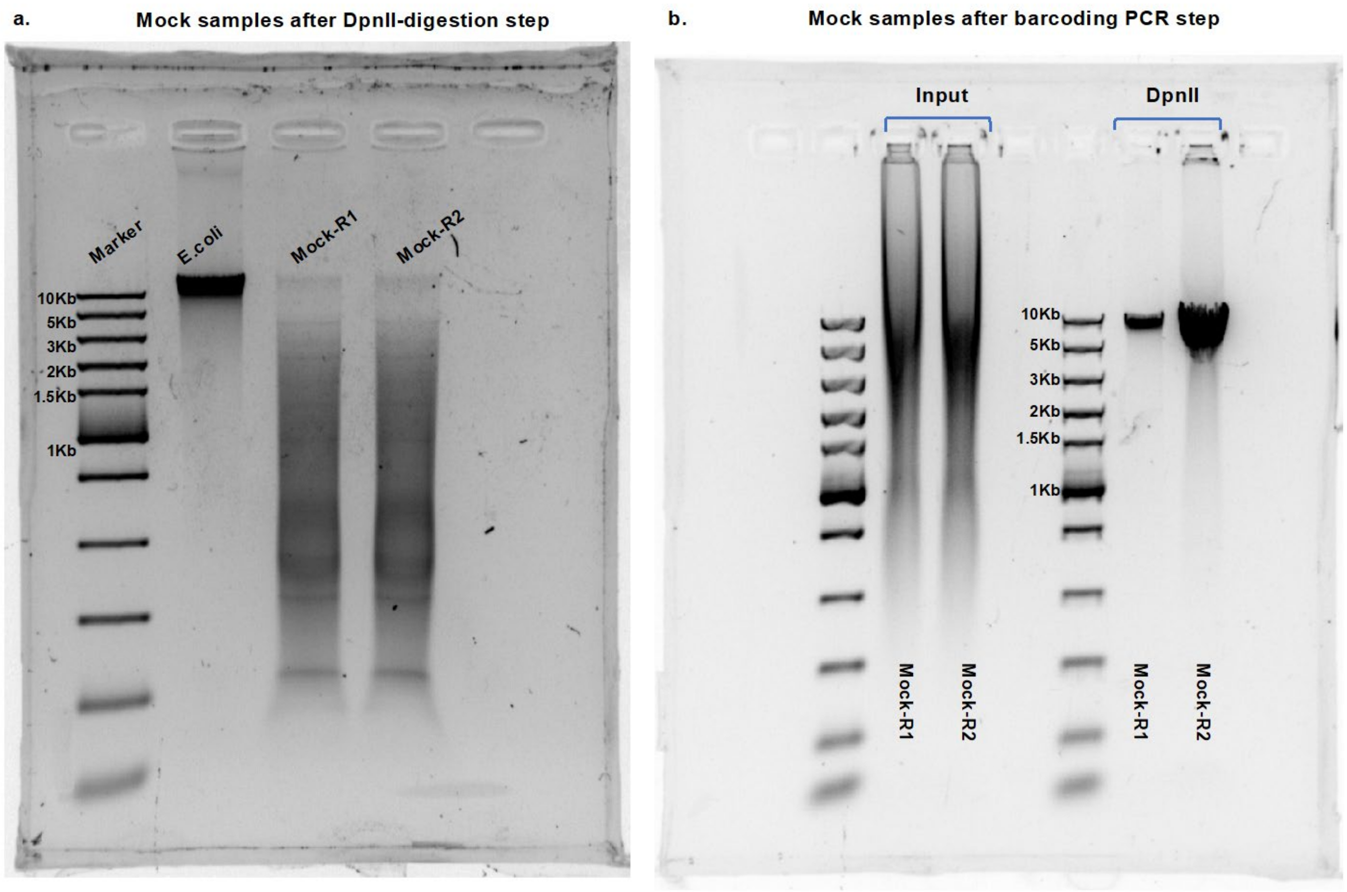


**Extended Fig. 4 Agarose gels of the two mock replicates. a**, Bands of ~10kb size are observed for two mock replicates digested by DpnII. *E. coli* DNA is used as the positive control for size reference in band collection from the gel (5- 10kb). **b,** The Input and DpnII-digested DNA were amplified by the barcoded primers and visualized on an agarose gel. The PCR bands for DpnII-digested DNA are 5- 10 kb size.

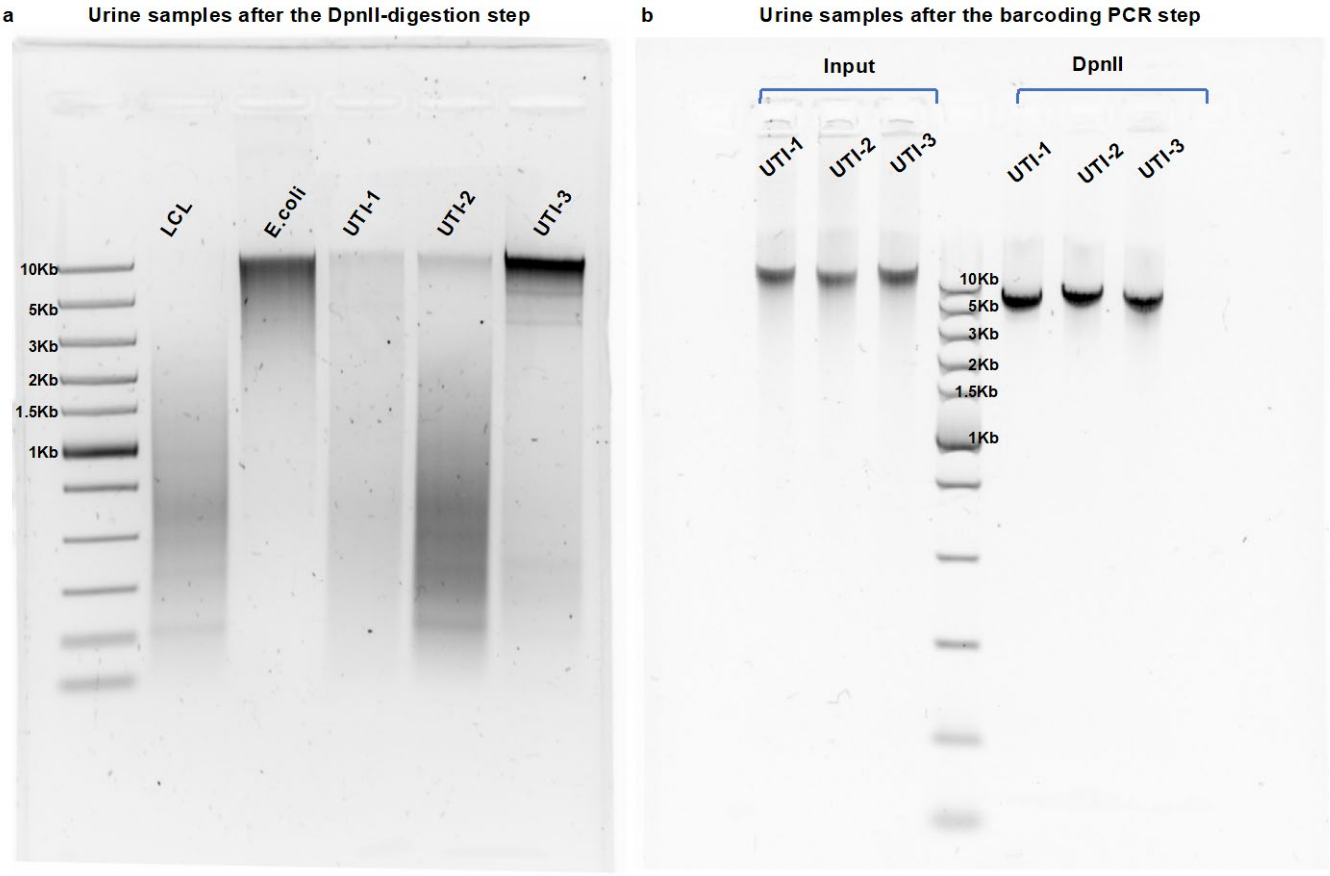


**Extended Fig. 5 Agarose gels for urine samples after the DpII-digestion and barcoding PCR steps.** **a,** RE-digested urine samples (UTI-1 to -3) are visualized on an agarose gel. Bands were observed, and cut out, for all three urine samples, around 5- 10kb in size. *E. coli* DNA (positive control) and lymphoblastoid cells (LCL, GM24149) DNA (negative control) are included to verify size estimations. **b,** A gel showing bands of the Input and DpnII-digested DNA that are amplified by the barcoded primers. PCR bands of DpnII-digested DNA are shown around 5- 10kb size.

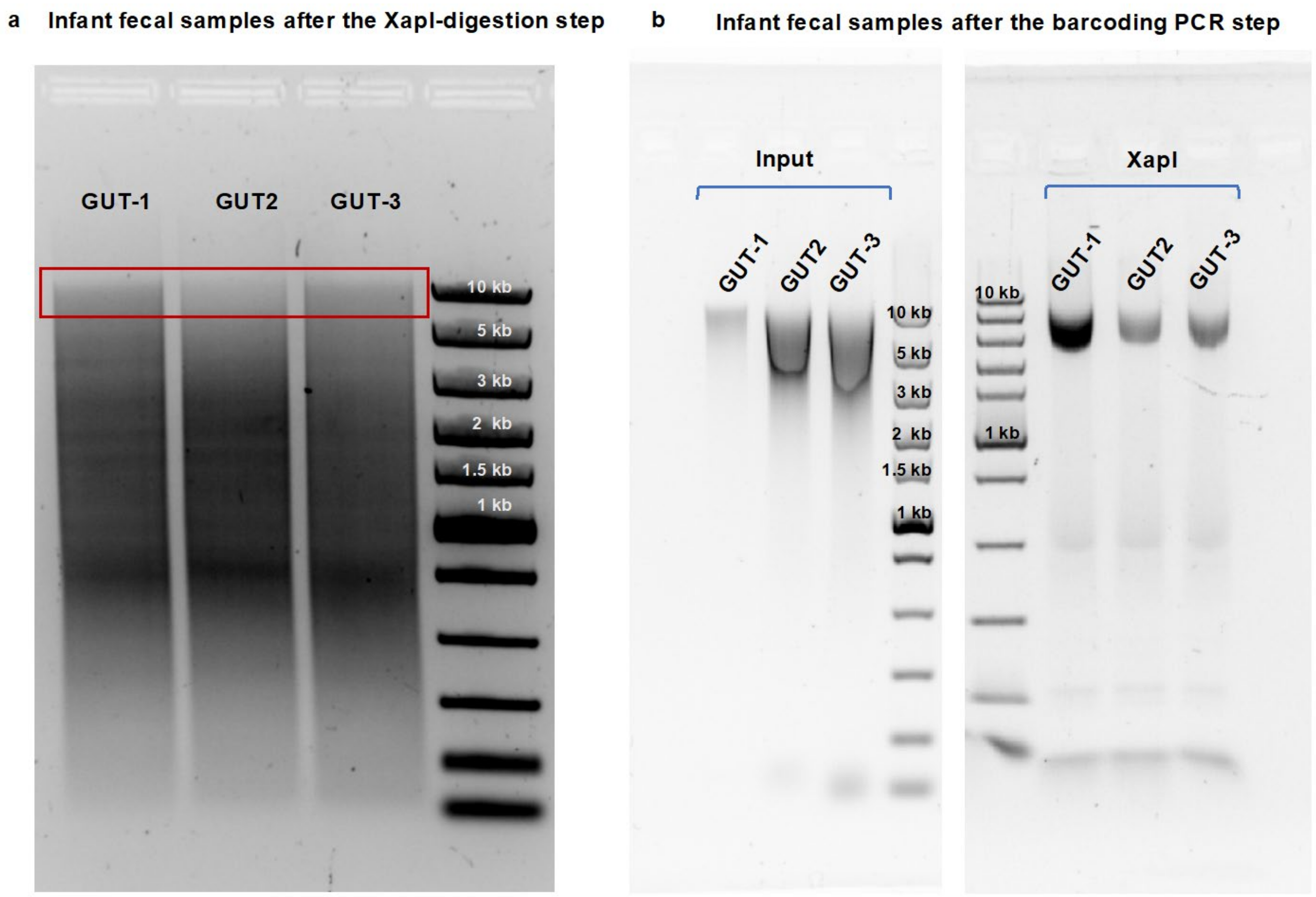


**Extended Fig. 6 Agarose gels for fecal samples after XapI-digestion and barcoding PCR steps. a,** A gel of the infant fecal samples (GUT-1 to -3) digested by XapI (recognition site: RAATTY). Faint (highlighted by the red square) bands are observed for the three fecal samples on an agarose gel. DNA was collected by cutting the gel at ~5- 10kb (based upon the DNA ladder). **b,** Gels show the bands of the Input (left panel) and XapI-digested DNA (right panel) that are amplified by the barcoded primers. The PCR bands of XapI-digested DNA are shown ~5- 10kb in size.

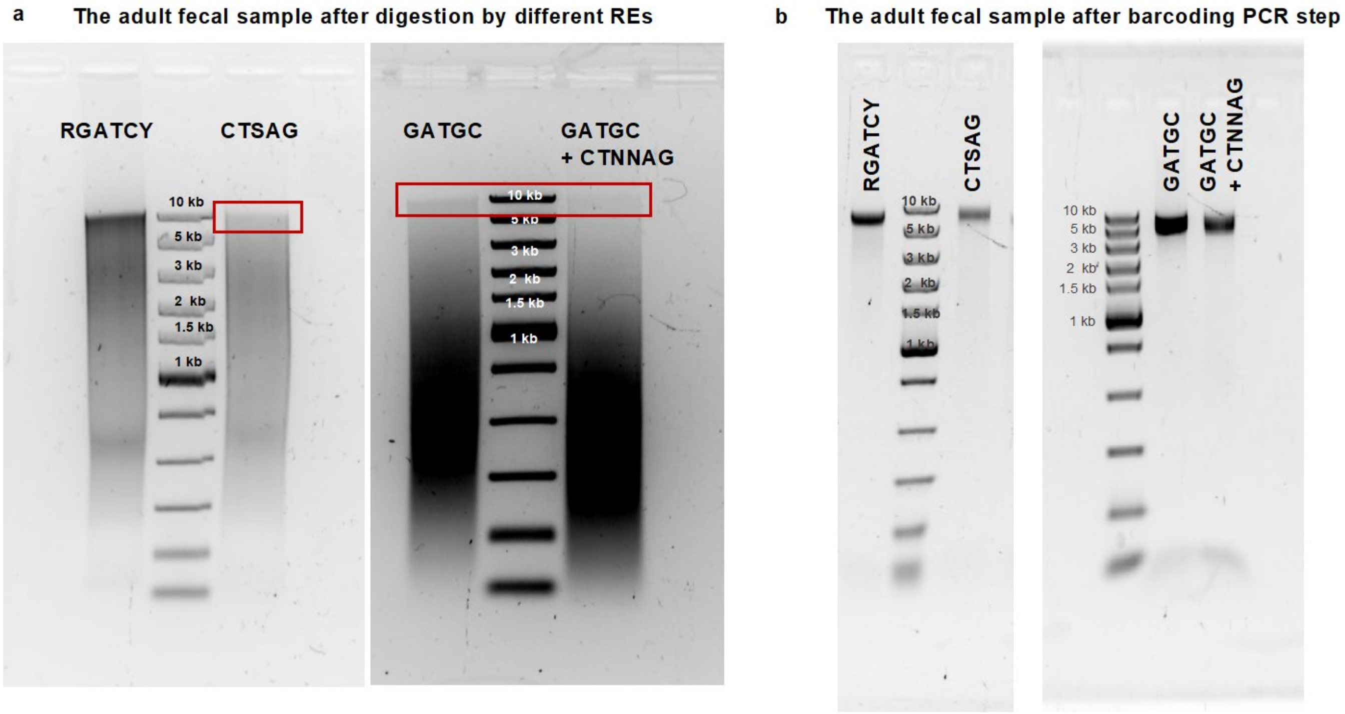


**Extended Fig. 7 Agarose gels of enriched low-abundance bacterial taxa from the adult fecal sample based on *de novo* discovered methylation motifs.** **a,** Gels for the adult fecal sample digested by different REs. Left, the MflI-digested adult fecal sample DNA (targeting RGATCY motif) shows the band at ~5- 10kb; the BspCNI (CTCAG as recognition site, part of the CTSAG motif)-digested sample shows a faint band at ~5- 10kb (based upon the DNA ladder). Right, faint bands (highlighted by the red square) are observed for the adult fecal sample digested by SfaNI (GATGC recognition site) and by the RE cocktails (targeting both GATGC and CTNNAG). **b,** Gels for PCR bands amplified by the barcoded primers, using RE-digested adult fecal DNA (from Fig. 7a) as the template. The PCR bands of RE(s)-digested DNA are shown around 5~10 kb.
